# Microevolution of *Pieris* butterfly genes involved in host-plant adaptation along a host-plant community cline

**DOI:** 10.1101/2021.09.30.462550

**Authors:** Yu Okamura, Ai Sato, Lina Kawaguchi, Atsushi J. Nagano, Masashi Murakami, Heiko Vogel, Juergen Kroymann

## Abstract

Herbivorous insects have evolved counteradaptations to overcome the chemical defenses of their host plants. Several of these counteradaptations have been elucidated at the molecular level, in particular for insects specialized on cruciferous host plants. While the importance of these counteradaptations for host plant colonization is well established,little is known about their microevolutionary dynamics in the field. In this study, we examine patterns of host plant use and insect counteradaptation in three *Pieris* butterfly species across Japan. The larvae of these butterflies express nitrile-specifier protein (NSP) and its paralog major allergen (MA) in their gut to overcome the highly diversified glucosinolate-myrosinase defense system of their cruciferous host plants. *Pieris napi* and *Pieris melete* colonize wild Brassicaceae whereas *Pieris rapae* typically uses cultivated *Brassica* as a host, regardless of the local composition of wild crucifers. As expected, NSP and MA diversity was independent of the local composition of wild Brassicaceae in *P. rapae*. In contrast, NSP diversity correlated with local host plant diversity in both species that preferred wild Brassicaceae. *P. melete* and *P. napi* both revealed two distinct major *NSP* alleles, which shaped diversity among local populations, albeit with different evolutionary trajectories. In comparison, *MA* showed no indication for local adaptation. Altogether, *MA* appeared to be evolutionary more conserved than *NSP*, suggesting that both genes play different roles in diverting host plant chemical defense.

## Introduction

Herbivorous insects encounter heterogeneous plant communities in the field and often use a subset of those plants as hosts. While feeding, herbivores are exposed to the chemical defenses of their host plants. Defenses typically vary between plant species and populations (Futuyma & Agrawal, 2009; Kliebenstein et al., 2001; Mitchell-Olds & Schmitt, 2006; Prasad et al., 2012; Windsor et al., 2005). Available evidence suggests that specialist herbivores acquired key innovations that enabled them to colonize their host plants by circumventing host plant chemical defenses (Berenbaum, Favret, & Schuler, 1996; Ratzka, Vogel, Kliebenstein, Mitchell-Olds, & Kroymann, 2002; Wheat et al., 2007; Wittstock et al., 2004). The microevolutionary dynamics of these herbivore counteradaptations is largely unknown, despite their ecological and evolutionary importance.

*Pieris* butterflies use plants from the family Brassicaceae as their hosts. These plants rely on the glucosinolate (GLS)-myrosinase system as their main chemical defense (Wheat et al., 2007; Wittstock et al., 2004; Wittstock & Halkier, 2002). GLSs are hydrolyzed by plant myrosinase enzymes when plant tissue is macerated by insect herbivory, and the main hydrolysis products, isothiocyanates, are highly toxic to most herbivores (Halkier & Gershenzon, 2006; Wittstock & Halkier, 2002). *Pieris* butterfly larvae overcome this defense system by expressing nitrile specifier proteins (NSPs) in their gut. NSPs affect the outcome of GLS breakdown to form nitriles rather than toxic isothiocyanates by an as yet unknown mechanism (Wittstock et al., 2004). Recent genetic and molecular analyses in *Anthocharis cardamines*, another Pierid species feeding on Brassicaceae, indicated that major allergen (MA), an ancient NSP paralog, has NSP-like activity (Edger et al., 2015). Noteworthy, *MA* transcript levels in the gut of *Pieris* larvae are upregulated in response to certain types of GLS (Okamura, Sato, Tsuzuki, Sawada, et al., 2019). Together, *NSP* and *MA* constitute the *NSP*-like gene family, a key innovation that enabled Pierid butterflies to colonize their host plants from the order Brassicales (Edger et al., 2015; Fischer, Wheat, Heckel, & Vogel, 2008; Wittstock et al., 2004).

GLSs are a highly diverse group of plant secondary metabolites, with about 135 different structures identified in the Brassicales (Blažević et al., 2020). GLS profiles vary both quantitatively and qualitatively (Agerbirk & Olsen, 2012; Fahey, Zalcmann, & Talalay, 2001; Olsen et al., 2016) and *Pieris* species typically use several genera within the Brassicaceae as their hosts (Friberg & Wiklund, 2019; Ohsaki & Sato, 1994; Ohsaki, 1979; Okamura, Sato, Tsuzuki, Sawada, et al., 2019; Okamura, Sawada, Hirai, & Murakami, 2016). Therefore, each species encounters a range of different GLS profiles in the field, a scenario that should result in local adaptation to host plant communities. Surprisingly, a previous study with *Pieris rapae*, the cabbage white butterfly, found no evidence for local adaptation of *NSP* genes in Europe or in the U.S. (Heidel-Fischer, Vogel, Heckel, & Wheat, 2010). However, this study did not address patterns of host-plant use. Furthermore, *P. rapae*, a considerable pest of *Brassica*, depends mainly on cultivated plants (Grishin et al., 2016; Ryan et al., 2019) and not on local populations of wild Brassicaceae. Hence, it seems important to test species that depend on wild plants in order to detect potential microevolutionary patterns in the genes that underlie insect counteradaptations against their host plants’ defenses.

In this study, we focus on three *Pieris* species, *Pieris melete*, *Pieris napi* and *P. rapae* that co-occur across most of Japan. *P. melete* and *P. napi* use wild Brassicaceae as host plants, including the genera *Arabis*, *Cardamine* and *Rorippa*, but their larvae are rarely observed on cultivated *Brassica* (Ohsaki & Sato, 1994; Okamura, Sato, Tsuzuki, Sawada, et al., 2019). In Japan, plant communities vary substantially along a north-south cline (Kubota, Shiono, & Kusumoto, 2015). Because this holds also true for the latitudinal distribution of wild Brassicaceae, we expected that the plant community cline should affect host-plant use in *P. melete* and *P. napi*, which depend on wild Brassicaceae, but not *P. rapae*, which feeds on cruciferous crops. We sampled local populations of these three species across Japan and acquired data about host plant communities at the sampling sites. We sequenced *NSP* and *MA* and used restriction-site associated DNA sequencing (RAD-seq) data for comparison, to distinguish between the potential impact of selection and neutral genetic processes (Bernatchez & Landry, 2003; Dionne, Miller, Dodson, Caron, & Bernatchez, 2007). As expected, we did not find any evidence for local adaptation of crop-dependent *P. rapae*. In contrast, we found a clear correlation between local host plant diversity and NSP, but not MA, amino-acid sequence diversity for both *Pieris* species that preferred wild Brassicaceae. Both *P. melete* and *P. napi* possessed two major *NSP* alleles, which caused the observed correlation with host plant diversity, albeit with completely different distribution patterns among Japanese populations. *P. napi* consisted of two distinct populations using very different sets of host plants, with a different *NSP* allele fixed in each population and evidence for directional selection of one allele in the past. In *P. melete*, on the other hand, two *NSP* alleles were maintained by balancing selection across Japan and allele frequency was correlated with local host plant diversity. Altogether, our study indicates that microevolution of counteradaptive traits has an important role for herbivores to adapt to heterogeneous chemical defenses in the host plants that they encounter in the field.

## Materials and Methods

### Selection of sampling sites

To determine suitable sampling sites, we estimated the diversity of Brassicaceae across Japan with Maxent ver. 3.4.1 (Phillips, Dudík, & Schapire, 2004; Phillips, Anderson, Dudík, Schapire, & Blair, 2017). We used 11,325 individual location data of 44 genera of the Brassicaceae in the Global Biodiversity Information Facility (GBIF.org; https://doi.org/10.15468/dl.a2nqtv) and S-Net (http://science-net.kahaku.go.jp), and 19 climate variables for Japan at 2.5 arc-minute resolution from WorldClim (Fick & Hijmans, 2017) to infer additional potential locations of genera from the Brassicaceae (Table S1). We calculated the Shannon index for genera-based diversity of Brassicaceae using the R package “vegan” (Oksanen et al., 2017). Based on this estimation, we chose eleven sampling sites across Japan for each of the three *Pieris* spp., *P. melete*, *P. napi* and *P. rapae* (Table S2).

**Table 1.**
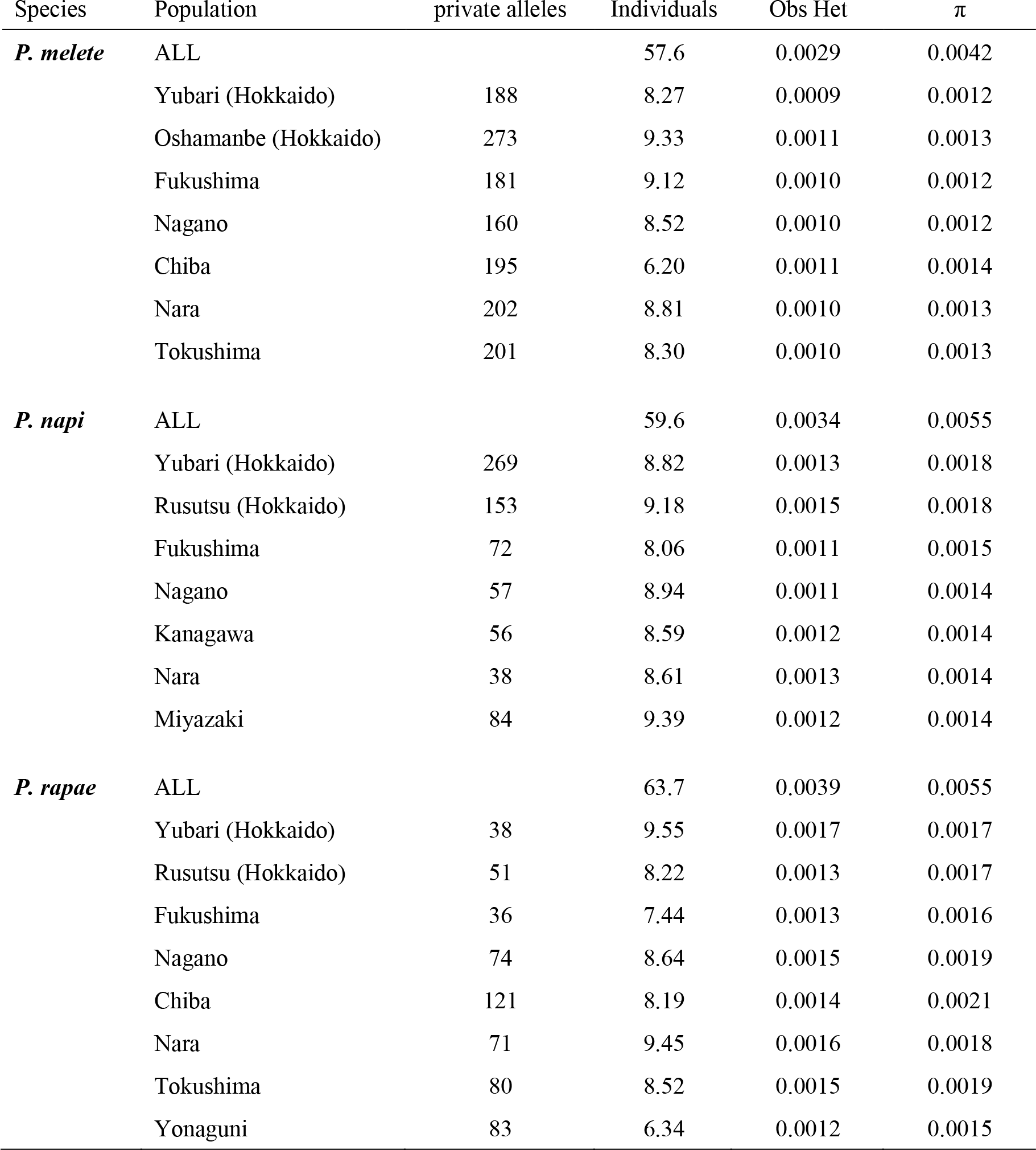
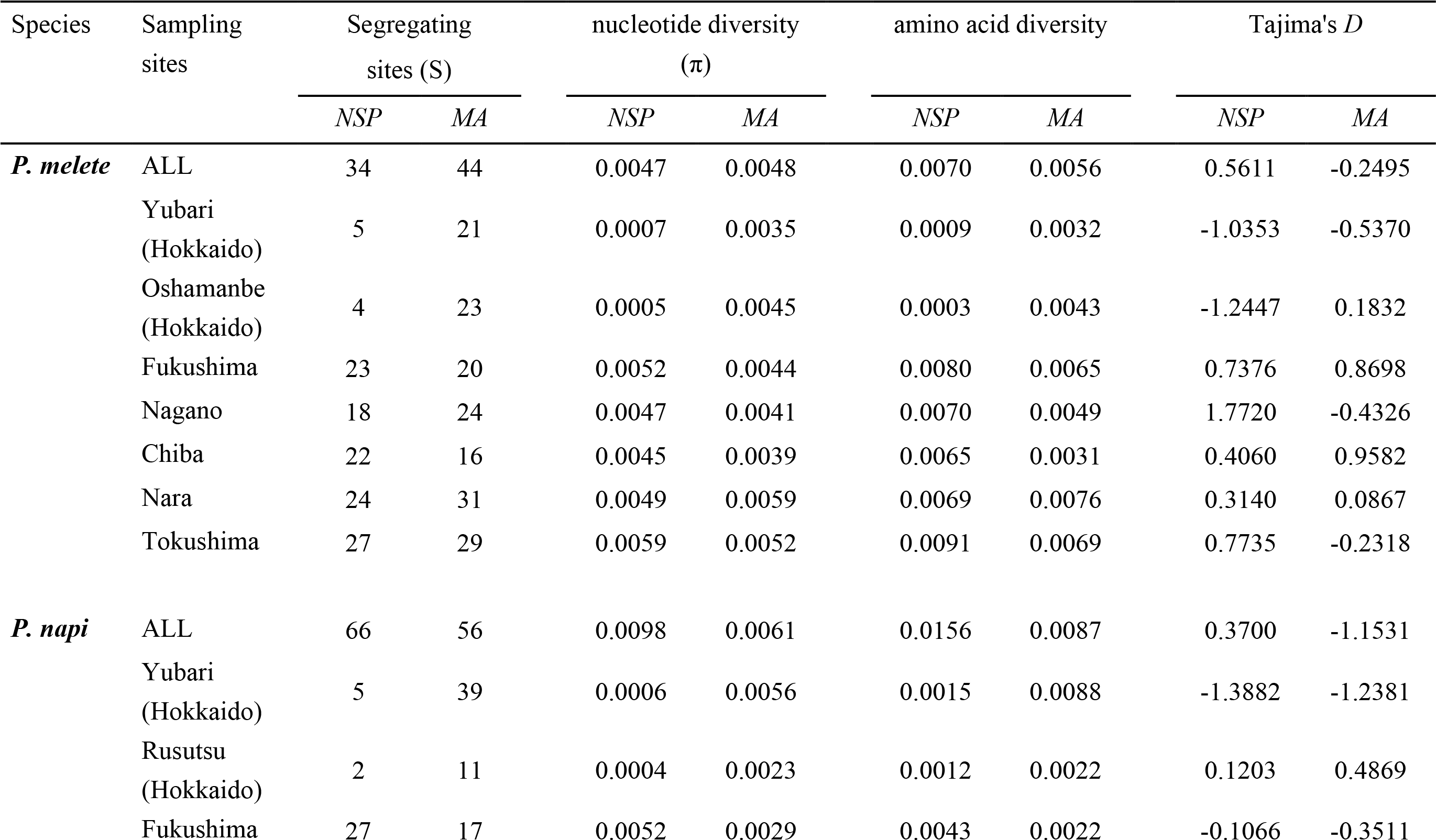
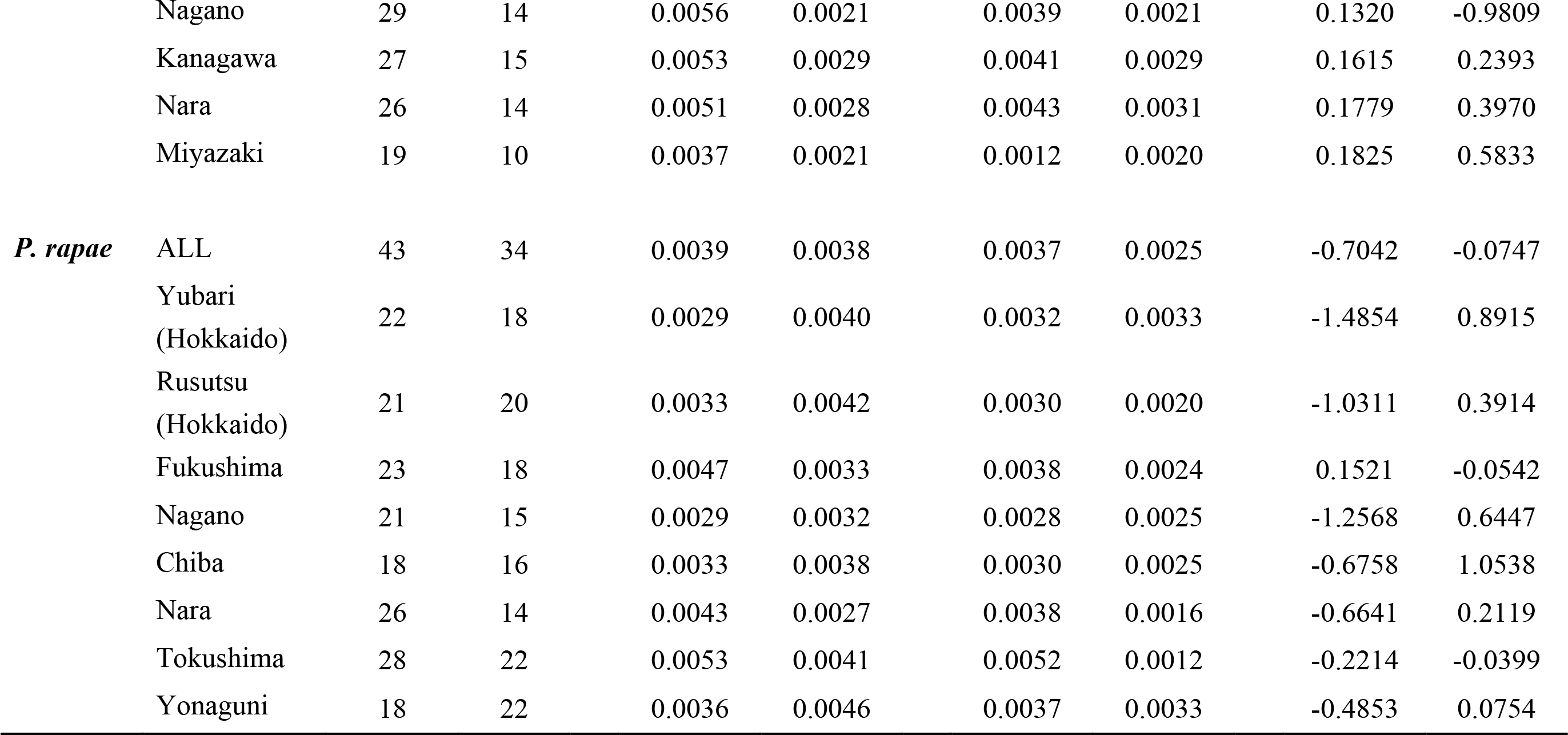
(a) Population genetic parameters from genome-wide RAD-seq in three Pieris Species (b) Population genetic parameters 702 for NSP and MA in three Pieris species

**Table 2.** Partial Mantel tests for NSP and MA in three *Pieris* species

| Comparison | Species | *r* | *P*-value |
| --- | --- | --- | --- |
| **Partial Mantel Test** |  |  |  |
| NSP AA Fst, Plant dissimilarity \| RAD Fst | *P. melete* | **0.444** | **0.039** |
|  | *P. napi* | **0.436** | **0.010** |
|  | *P. rapae* | -0.047 | 0.594 |
| MA AA Fst, Plant dissimilarity \| RAD Fst | *P. melete* | 0.1405 | 0.258 |
|  | *P. napi* | -0.318 | 0.867 |
|  | *P. rapae* | -0.183 | 0.877 |

NSP AA Fst and MA AA Fst: Fst-like divergences based on amino acid sequences. Plant dissimilarity: Bray-Curtis plant community dissimilarity. Statistically significant results are in bold ( $P \leq 0.05$ ).

### Sampling of Brassicaceae and *Pieris* spp

We collected *Pieris* larvae from April to August 2017. At each sampling site, we used the transect method to search for Brassicaceae along a > 2-km sampling path. We noted the number of plants per species, and collected larvae or eggs of *P. melete*, *P. napi* and *P. rapae* from their host plants. We disregarded two Brassicaceae genera, *Capsella* and *Erysimum*, because these are unsuitable hosts for those *Pieris* species (Okamura et al., 2016). Similarly, we did not take into account cultivated Brassicaceae, *i.e.*, cabbage and kale, growing in crop fields near sampling sites. We calculated the host plant diversity using Shannon diversity index for each sampling site, based on the number of species and plants per species. We provided eggs or small larvae with their host plants until they reached the 3^rd^ instar and then dissected the larvae to separate guts from the rest of the bodies. We processed larvae larger than 3^rd^ instar immediately. We stored guts in RNAlater (QIAGEN) and body rests at -80°C. We excluded larvae that contained parasitoid wasps from further analyses.

### RFLP-based identification of *Pieris* butterfly species

We used restriction fragment length polymorphisms (RFLPs) in the mitochondrial *ND5* gene (GenBank accession numbers: LC090587- LC090590) to determine species identity of larvae. *P. melete ND5* has a *Hin*cII site, and *P. napi ND5* has a *Hin*fI and a *Hin*dIII site, whereas *P. rapae ND5* lacks these restriction sites. We amplified *ND5* directly from larval bodies, using MightyAmp™ DNA Polymerase Ver.3 (TaKaRa) and *ND5* universal primers (Table S3), followed by restriction and separation on 2% agarose gels. To verify that our species assignments worked correctly, we used the same RFLP technique for adult males previously classified as *P. melete* (N = 24), *P. napi* (N = 16) or *P. rapae* (N = 24) based on differences in the shape of the androconium. *ND5*-based identification matched androconium-based identification for all 64 individuals.

**Table 3.**
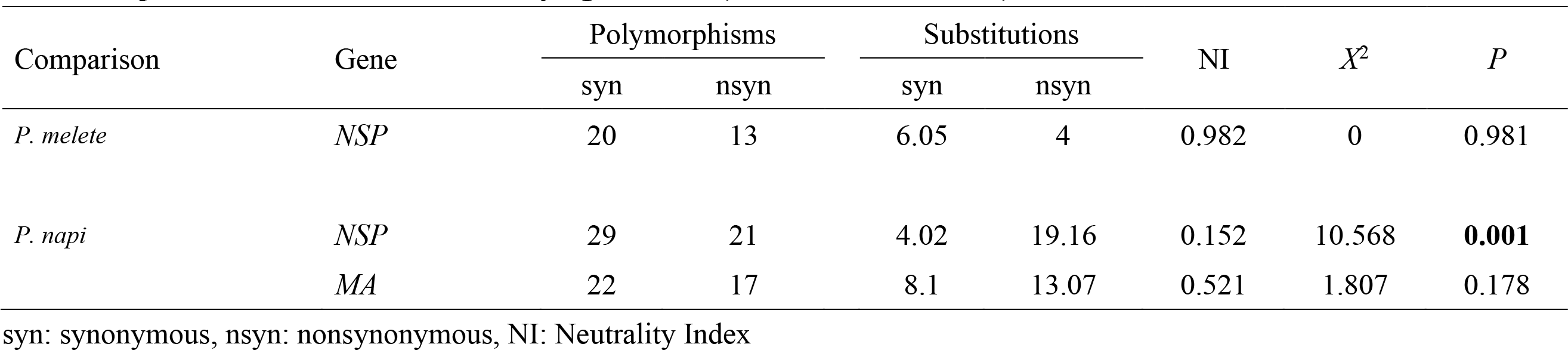
Population-based tests for diversifying selection (McDonald–Kreitman)

### *NSP* and *MA* amplification, cloning and sequencing

We randomly picked ten larvae per species and sampling site to sequence *NSP* and *MA* genes. In total, we used 70 larvae (ten larvae per site from seven sampling sites) of *P. melete* and *P. napi,* respectively, and 78 larvae of *P. rapae* (ten larvae per site from seven sampling sites, plus eight larvae from Yonaguni Island). We extracted RNA from dissected larval gut using the RNeasy Mini Kit (QIAGEN), synthesized cDNA with the ReverTra Ace qPCR RT Master Mix (TOYOBO) and used TaKaRa Ex Taq (TaKaRa) for amplification of *NSP* and *MA* genes. We gel-purified PCR products with the GEL/PCR Purification Mini Kit (FAVORGEN). We designed PCR primers for *NSP* and *MA* genes based on available RNA-seq data (Accession numbers: ERX2829492- ERX2829499) to confirm that primer-binding sites were conserved and to avoid a potential allelic bias in amplification (Table S3). We cloned it with Mighty TA-cloning kit (TaKaRa), purified plasmids with NucleoSpin Plasmid EasyPure (TaKaRa), and sequenced one plasmid for each *NSP* and *MA* gene from each of the 218 larval samples, using an ABI 3730xl DNA Analyzer (Applied Biosystems).

### Network analysis of *NSP* and *MA*, and PCR-RFLP of *P. melete NSP* gene variants

We trimmed and aligned *NSP* and *MA* reads with MEGA6 (Tamura, Stecher, Peterson, Filipski, & Kumar, 2013) and Mafft (Katoh & Standley, 2013). For each species, we excluded singleton SNPs, assuming that they were PCR errors. We performed NeighborNet network analyses with SplitsTree 4.15.1 (Huson & Bryant, 2006). We used PCR-RFLP to distinguish two major *NSP* variants in *P. melete* to determine whether these variants were different gene copies or alleles. We amplified *NSP* exon 1 from gDNA with EmeraldAmp MAX PCR Master Mix (TaKaRa), using primers specific for exon 1 (Table S3), followed by a *Hin*cII digest to target a fixed SNP, and separation on 3% TBE agarose gels.

### RT-qPCR of *P. melete NSP* and *MA* genes

To determine whether *NSP* and *MA* gene copy numbers differed among *P. melete* genotypes, we conducted Real Time quantitative PCR (RT-qPCR) from gDNA using *Ef1a* as a reference gene. For primer design, we used Primer3Plus (Untergasser et al., 2007) with a product size of 70 to180 bp, a Tm of 59 to 61°C, a GC content of 40 to 60% and a maximum polybase of 3. For RT-qPCR, we used the CFX Connect Real-Time PCR Detection System (Bio-Rad) using TB Green Premix Ex Taq II (Tli RnaseH Plus) (TaKaRa). We used the ddCq method (Pfaffl, 2001) and ANOVA to compare relative gene copy numbers among genotypes.

### RAD sequencing

We extracted gDNA for RAD-seq from the same 218 individuals that we had used for sequencing of *NSP* and *MA* genes, using the Maxwell 16 LEV Plant DNA Kit (Promega), with *Eco*R1 as the restriction enzyme for generating RAD-seq libraries. We ran all 218 samples in a single lane on a HiSeq2500 (Illumina SE 50 bp) and trimmed reads with trimmomatic (Bolger, Lohse, & Usadel, 2014), using ILLUMINACLIP:2:10:10, TRAILING:20, SLIDINGWINDOW:4:15, and MINLEN:30. We excluded samples with less than 500,000 reads from further analysis. We called SNPs with stacks ver. 1.48 (Catchen, 2013). For ustacks, we set n = 3 and M = 3, and for cstacks, we used the n = 3 option. We performed this analysis for each species independently. To acquire values for genetic diversity (π) and genetic distance (Fst) of each population, we used “populations” with parameter p set as sampled population number and r = 0.70. Since we suspected that *P. melete* and *P. napi* could hybridize, we also called SNPs by setting p = 1 and r = 0.85 in multiple species scales without involving population information for assessing genetic structure of the three species.

### Population and evolutionary genetic analyses

For analyses of potential population structure we used RAD-seq data in multiple species scales and performed sparse non-negative matrix factorization (SNMF) with the “snmf” function (*K* = 1 to 10, repetition = 20, iterations = 200) implemented in the R package “LEA” (Frichot & François, 2015; Frichot, Mathieu, Trouillon, Bouchard, & François, 2014). We used cross-entropy criterion to determine the number of ancestral population (*K*). We calculated species-wide distributions of Tajima’s *D* (Tajima, 1989) of *NSP* and *MA* using the poly-div_sfs.pl script (https://github.com/santiagosnchez/poly-div_sfs) iterating random sampling of one individual per population and calculation of *D* for 300 times. We performed these analyses on both, entire coding sequence and individual exons. For calculation of genome-wide Tajima’s *D* at the species level, we used VCFtools (Danecek et al., 2011). We did the same random sampling coupling with selecting 30 random SNPs corresponding to the average number of SNPs in *NSP* within populations. We used poly-div_sfs.pl script to calculate allele frequency of *NSP* and *MA* for each species. At the population level, we used DNAsp ver.6 (Rozas et al., 2017) for calculation of π and Fst of *NSP* and *MA*, and Arlequin ver. 3.5 (Excoffier & Lischer, 2010) for Tajima’s *D*. We estimated Fst-like amino acid divergence in NSP and MA between populations as follows; Fst_AA = {*Dij* – mean (*D* within population)} / *Dij*, where *Dij* shows mean p-distance in amino acid sequences between population *i* and *j*. We used the online tool at mkt.uab.es/mkt/MKT.asp (Egea, Casillas, & Barbadilla, 2008) to conduct McDonald–Kreitman tests (McDonald & Kreitman, 1991).

### Comparison of *NSP* and *MA* with host-plant diversity

We used linear regression to test for each *Pieris* species whether diversity of *NSP* or *MA* correlated with host-plant diversity across sampling sites, using measured host plant diversity as the explanatory variable and NSP or MA amino acid diversity, or genome-wide genetic diversity estimated from RAD-seq data as response variables. Furthermore, we performed a partial Mantel test (Mantel, 1967; Smouse, Long, & Sokal, 1986; Sokal, 1979) to test whether NSP or MA amino acid sequence divergence between *Pieris* populations correlated with host-plant community dissimilarities, using the “vegan” package in R (Oksanen et al., 2017). We used the partial Mantel test to compare Fst-like amino acid sequence distances of NSP or MA with the Bray-Curtis index for host plant dissimilarity, after controlling for genetic differences between populations (Fst) using RAD-seq data. We permuted rows and columns of the first dissimilarity matrix 100,000 times to evaluate the reliability of the partial Mantel test results.

## Results

### 1. Field Sampling

#### Diversity of Brassicaceae in the field reflects Maxent-based predictions

The Maxent presence-probability estimation predicted that diversity in Brassicaceae communities should be higher in central Japan than in northern and southern regions (Fig. 1a). We chose eleven sites along this predicted cline of Brassicaceae diversity: Yubari, Rusutsu and Oshamanbe (all: on the island of Hokkaido), Fukushima, Nagano, Chiba, Kanagawa, Nara (all: on the island of Honshu), Tokushima (on the island of Shikoku), Miyazaki (on the island of Kyushu), and the island Yonaguni (from north to south; Fig. 1a). In total, we collected data on 4,777 individual plants across 14 genera and 25 species of the Brassicaceae (Table S2). We used these data to estimate the Shannon index for the diversity of Brassicaceae communities at each site. As expected, field-observed diversity in Brassicaceae communities correlated positively with Maxent predictions (*r* = 0.701, *P* = 0.016), was highest in central Japan, and declined towards the north and the south (Fig. 1b).

**Fig. 1.**
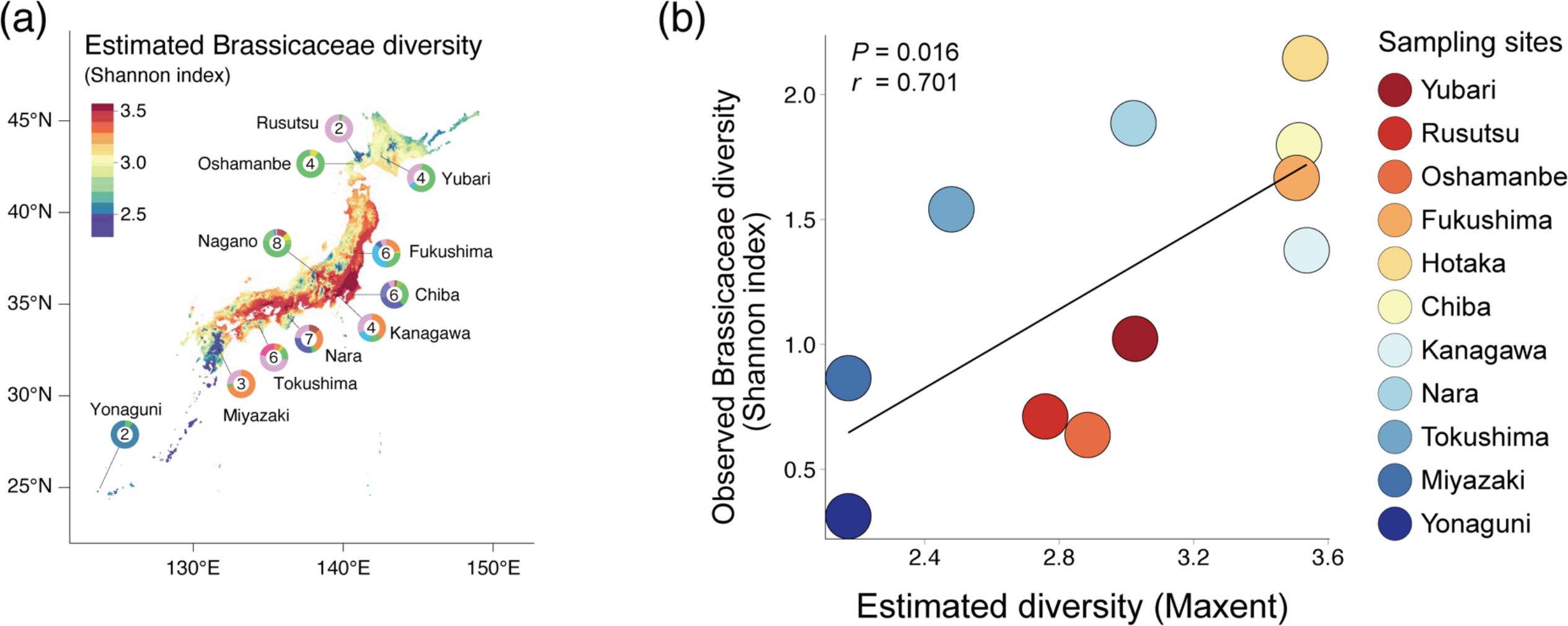
(a) Maxent-based diversity in Brassicaceae across Japan. The color code indicates the Shannon index for estimated diversity. Pie charts show number and composition of observed Brassicaceae genera at each sampling site. (b) Pearson correlation between estimated and observed diversity of Brassicaceae at each sampling site, with the north-south order of sampling sites depicted to the right of the panel. Diversity in the Brassicaceae is higher in central Japan than in northern or southern Japan.

#### Sampling of *Pieris* larvae in the field

In total, we collected 945 larvae from eleven sampling sites (Table S4). Based on PCR-RFLP using restriction site polymorphisms in the mitochondrial *ND5* region, we identified 483 larvae of *P. melete*, 253 of *P. napi* and 209 of *P. rapae*.

### 2. Population structure of three *Pieris* species in Japan

We randomly chose ten individual larvae per population and species for RAD-seq. On average, we obtained 1,062,550 RAD-seq reads per sample. After removal of samples with low read counts (< 500,000 reads), 183 samples remained for population genetic analyses. Stacks ver.1.48 called 2,490 SNPs for *P. melete*, 1,376 for *P. napi*, and 1,410 for *P. rapae.* Genome-wide diversity was higher in *P. rapae* and *P. napi* (both: π = 0.055) than in *P. melete* (π = 0.042) (Table 1a). Individual populations of *P. melete* displayed relatively uniform diversity among sampling sites. *P. rapae* populations from central Japan tended to have slightly higher diversity than northern or southern populations. Finally, Hokkaido populations of *P. napi* had higher diversity than other populations in the remainder of Japan (Table 1a)

Genome-wide data did not suggest any apparent population genetic structure of *P. melete* or *P. rapae* across Japan, in contrast to *P. napi* (Fig. 2, 3d). In *P. napi*, populations from Hokkaido were distinct from populations in other parts of Japan. In addition, we found evidence for some, very limited gene flow between species (Fig. 2) but not for recent hybridization events among these closely related butterfly species.

**Fig. 2.**
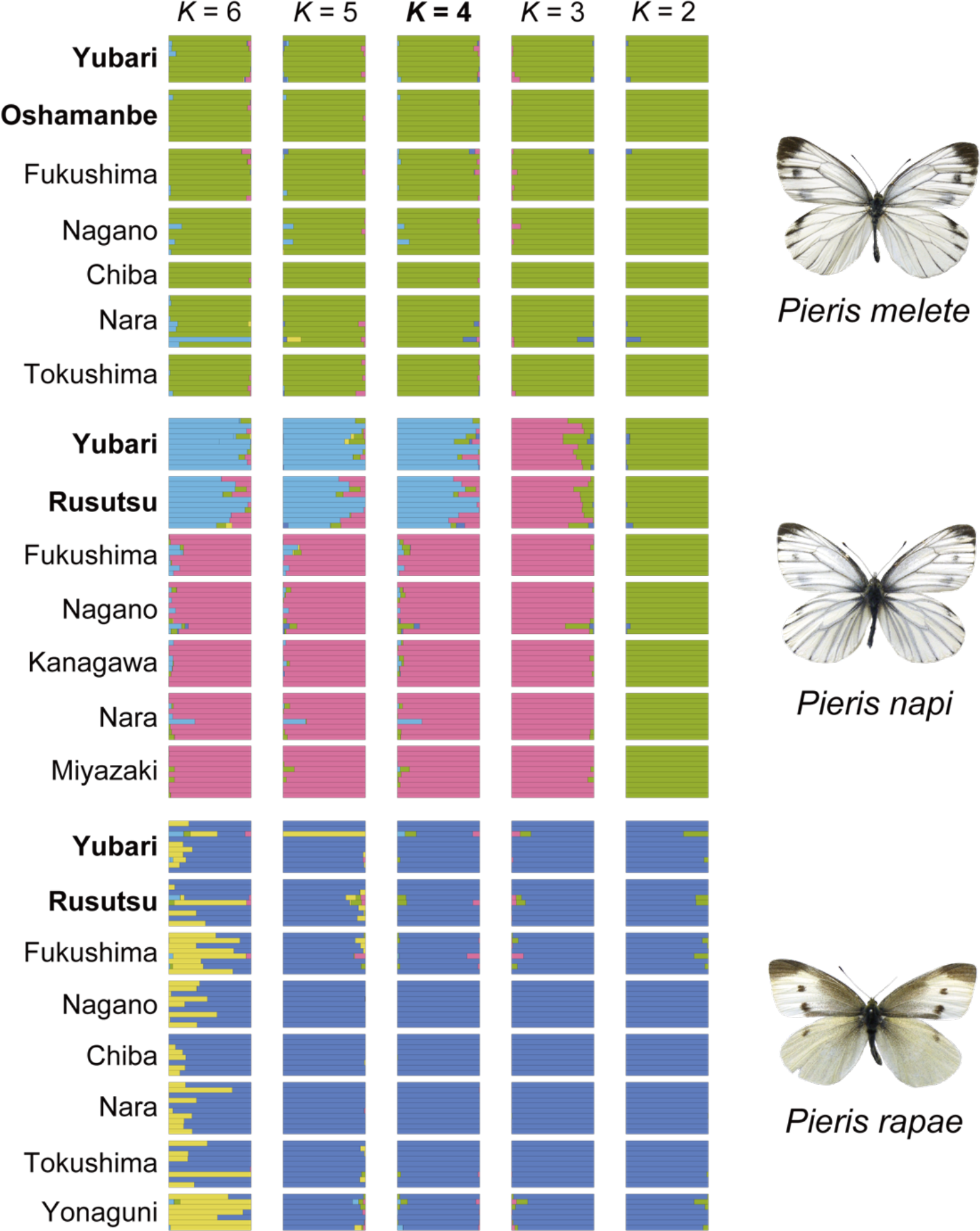
Inferred population genetic structures of three *Pieris* spp. across Japan for different clustering numbers (*K*), based on RAD-seq data. The cluster number most highly supported by the cross-entropy criterion (*K* = 4) is shown in bold. Colored horizontal lines correspond to individual samples, and blocks correspond to a population at a particular sampling site, ordered from north to south with Hokkaido sites in bold.

**Fig. 3.**
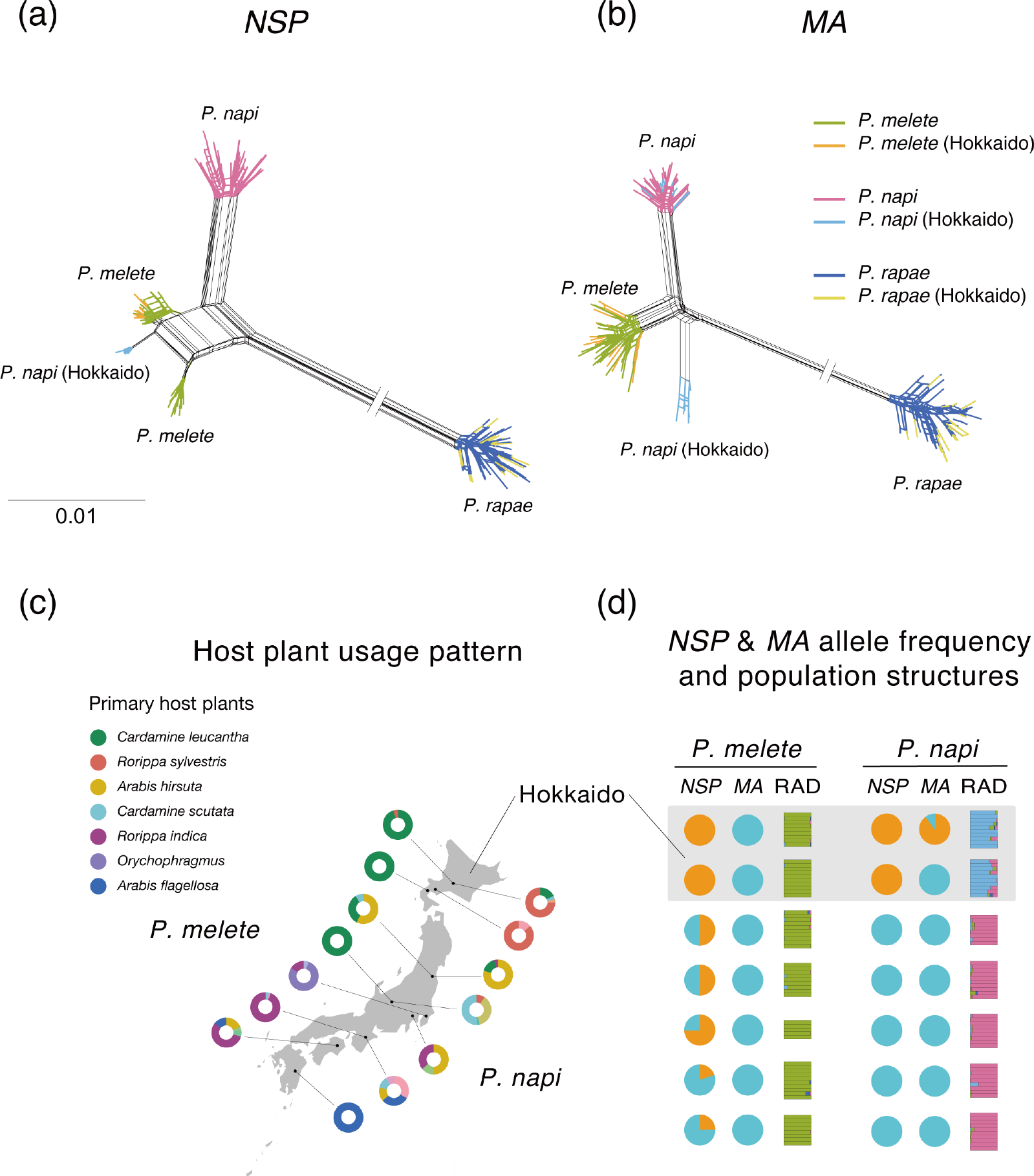
NeighborNet networks of (a) *NSP*s and (b) *MA*s from three *Pieris* species across Japan, with different colors for species and for Hokkaido populations. Note that *P. melete* and *P. napi* both have two major *NSP* alleles. Note further that *P. napi* has two major *MA* alleles. (c) Observed host plant usage patterns of *P. melete* and *P. napi* at sampling sites. (d) Allele frequencies of major *NSP* and *MA* alleles in comparison with genome-wide genetic structure of *P. melete* and *P. napi* populations at each sampling site, ordered from north to south.

### 3. Diversity of *NSP* and *MA*

#### *NSP* and *MA* sequences are diverse but single copy in *P. melete* and *P. napi*

We cloned and sequenced *NSP* and *MA* cDNAs from the same larvae that we used for RAD-seq. PCR products contained near full-length coding sequences (CDS) of *NSP* (1,860 bp of a total CDS of 1,896 bp) and *MA* (1,869 bp of 1,899 bp).

Network analysis identified two main clusters of *NSP* variants in both, *P. melete* and *P. napi* (Fig. 3a). In contrast, all *P. rapae NSP* sequences grouped together, separate from *P. melete* and *P. napi NSP*s. Hokkaido populations of *P. melete* were fixed for one of the main *NSP* variants (Figs. 3a, d), while all other *P. melete* populations possessed both variants, albeit with an overall decline in the frequency of the Hokkaido variant towards the south (Fig. 3d). One of the *P. napi NSP* variants was only found in populations from Hokkaido, while the other variant was fixed in central and southern populations (Fig. 3a, d). Network analysis of *MA* showed one single sequence cluster for each of *P. melete* and*P. rapae,* while *P. napi* sequences fell into two distinct clusters (Fig. 3b), with one of the two *P. napi MA* variants present only in Yubari on Hokkaido (Figs. 3b, d).

To test whether the different *NSP* variants of *P. melete* corresponded to different alleles or two different gene copies, we conducted two experiments, a discrimination of variants with PCR-RFLP and a RT-qPCR-based quantification of gene copy number. The *P. melete NSP* variants differed by a SNP causing a *Hin*cII restriction site polymorphism. After PCR, restriction digest, and agarose gel electrophoresis, we expected to obtain one undigested and one digested PCR product across all individuals if *NSP* variants corresponded to two distinct gene copies. In contrast, we expected to obtain undigested and digested PCR products in various combinations in the case that *NSP* variants corresponded to segregating alleles of the same gene. Indeed, we found samples with only undigested or with only digested PCR products, as well as samples with a mix of both, indicating that *NSP* variants corresponded to different alleles of the same gene (Fig. S1). Furthermore, gene copy numbers determined by RT-qPCR (Fig. S2) from eight randomly chosen samples per genotype were compatible with the same number of *NSP* copies in all *P. melete* genotypes, confirming that *NSP* variants were indeed alleles of the same gene.

### 4. NSP and MA diversity and plant community diversity

#### NSP but not MA amino acid diversity correlates positively with host-plant diversity in *P. melete* and *P. napi*

We found larvae of *P. melete* feeding on eleven, *P. napi* on 14, and *P. rapae* on nine different plant species (Table S4). We found 36% of all *P. melete* larvae on *Rorippa indica*, 19% on *Cardamine leucantha*, 18% on *Orychophragmus violaceus*, and 13% on *Arabis hirsuta* (Table S4). *P. napi* larvae were mostly present on *Rorippa sylvestris* (23%), *Arabis flagellosa* (17%), *Cardamine occulta* (17%), *Arabis hirsuta* (12%) and *Barbarea orthocera* (11%). Finally, we found *P. rapae* larvae most often on feral plants of *Brassica napus* (33%), followed by *Rorippa indica* and *Rorippa sylvestris* (both: 15%), *Cardamine kiusiana* (12%), *Nasturtium officinale* and *Rorippa palustris* (both: 10%). However, we also saw a substantial number of *P. rapae* larvae and eggs in crop fields of cabbage, broccoli or kale near sampling sites but we did not have permission to sample from these fields.

Altogether, *Pieris* species appeared to use a more diversified set of host plants in central Japan than in northern and southern Japan (Fig. 3c, Table S4). Therefore, we compared diversity in detoxification genes with the diversity of Brassicaceae at sampling sites (Fig. 4a). Local host plant diversity correlated significantly with NSP amino acid diversity for both *P. melete* (*r* = 0.83, *P* = 0.022) and *P. napi* (*r* = 0.88, *P* = 0.010), but not *P. rapae* (*r* = 0.021, *P* = 0.961). By contrast, local host plant diversity did not correlate with MA amino acid diversity or genome-wide diversity for any of the three species (Fig. 4b).

**Fig. 4.**
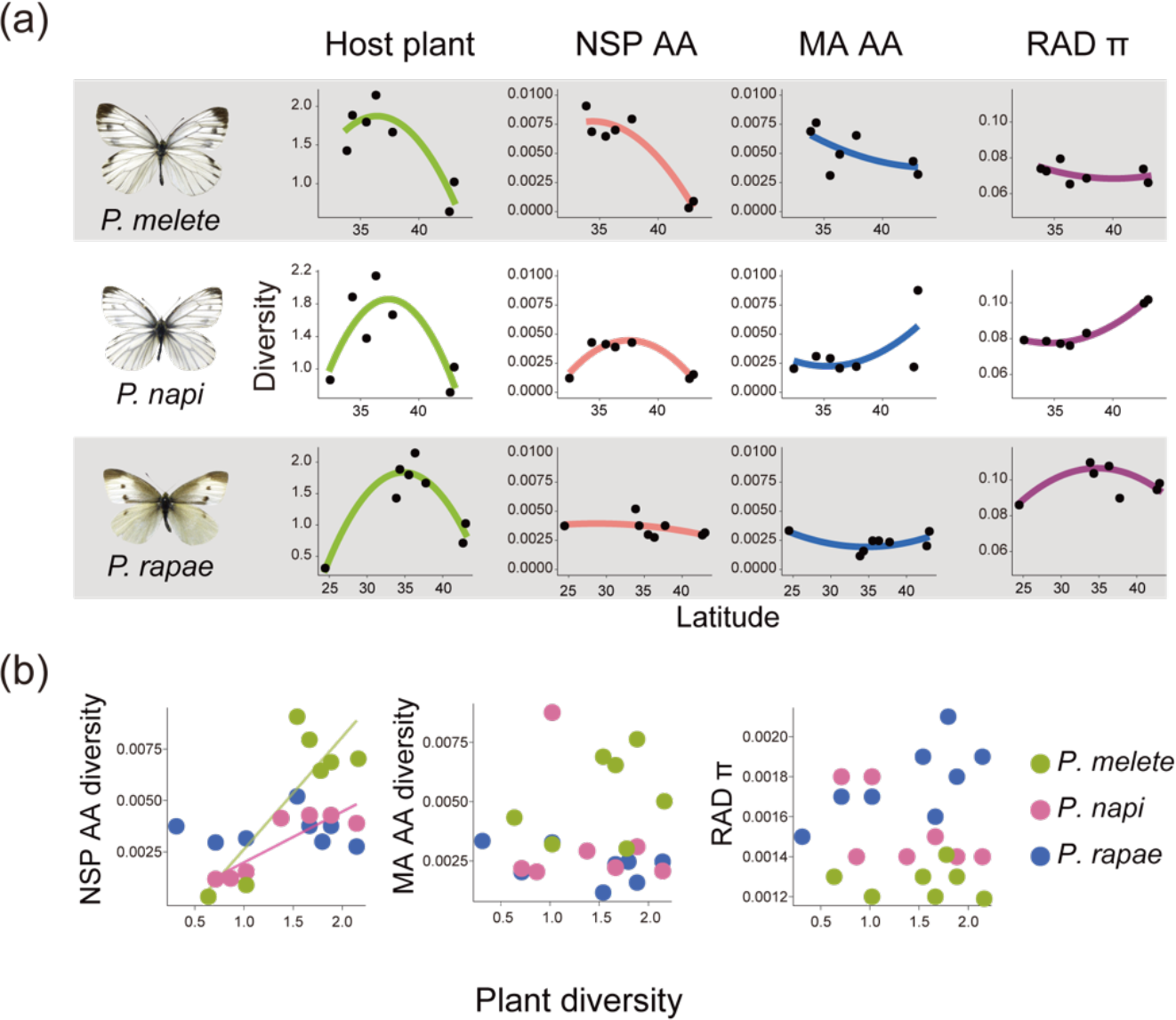
(a) Host plant diversity (Shannon index), NSP amino acid diversity (NSP AA), MA amino acid diversity (MA AA), and genome-wide diversity (RAD π) within three *Pieris* species at sampling sites ordered according to latitude. Black dots correspond to sampling sites, with colored lines showing regression curves. (b) Correlations between NSP AA, MA AA, or RAD π with host plant diversity in three *Pieris* species. Note the significant positive correlation (*P* < 0.05) between plant diversity and NSP AA diversity in *P. melete* and *P. napi*, but not in *P. rapae*.

To confirm the relationship between host plant and NSP sequence, we conducted a partial Mantel test (Mantel, 1967; Smouse et al., 1986; Sokal, 1979). Again, we found positive correlations between host plant dissimilarities and NSP sequence divergence between populations for both, *P. melete* (*r* = 0.443, *P* = 0.039) and *P. napi* (*r* = 0.436, *P* = 0.009) but not *P. rapae*. Likewise, results for MA remained non-significant (Table 2).

### 5. Statistical tests for selection

#### Different evolutionary regime between *NSP* and *MA*

To identify patterns of potential selection, we calculated Tajima’s *D* for *NSP* (*DNSP*), *MA* (*DMA*), and at the genome-wide level (*DRAD*). *DRAD* was negative in all three species, compatible with overall population expansion (Fig. 5a). In contrast, *DNSP* was positive and substantially higher than *DRAD* in *P. melete* and *P. napi*. Also, *DMA* was higher than *DRAD* in *P. rapae*. Furthermore, *DNSP* varied substantially among exons in *P. melete* and moderately in *P. napi* but not in *P. rapae*, while *DMA* displayed a more uniform pattern across exons of all three species (Fig. 5b). In *P. melete*, exon 1 had a particularly high positive *D*. Similarly, *P. napi* exon 3 displayed an elevated *D*. Close inspection of these exons revealed that both contained several linked, nonsynonymous SNPs at intermediate frequency (Fig. 5c), suggesting that *NSP* was under balancing selection in *P. melete*.

**Fig. 5.**
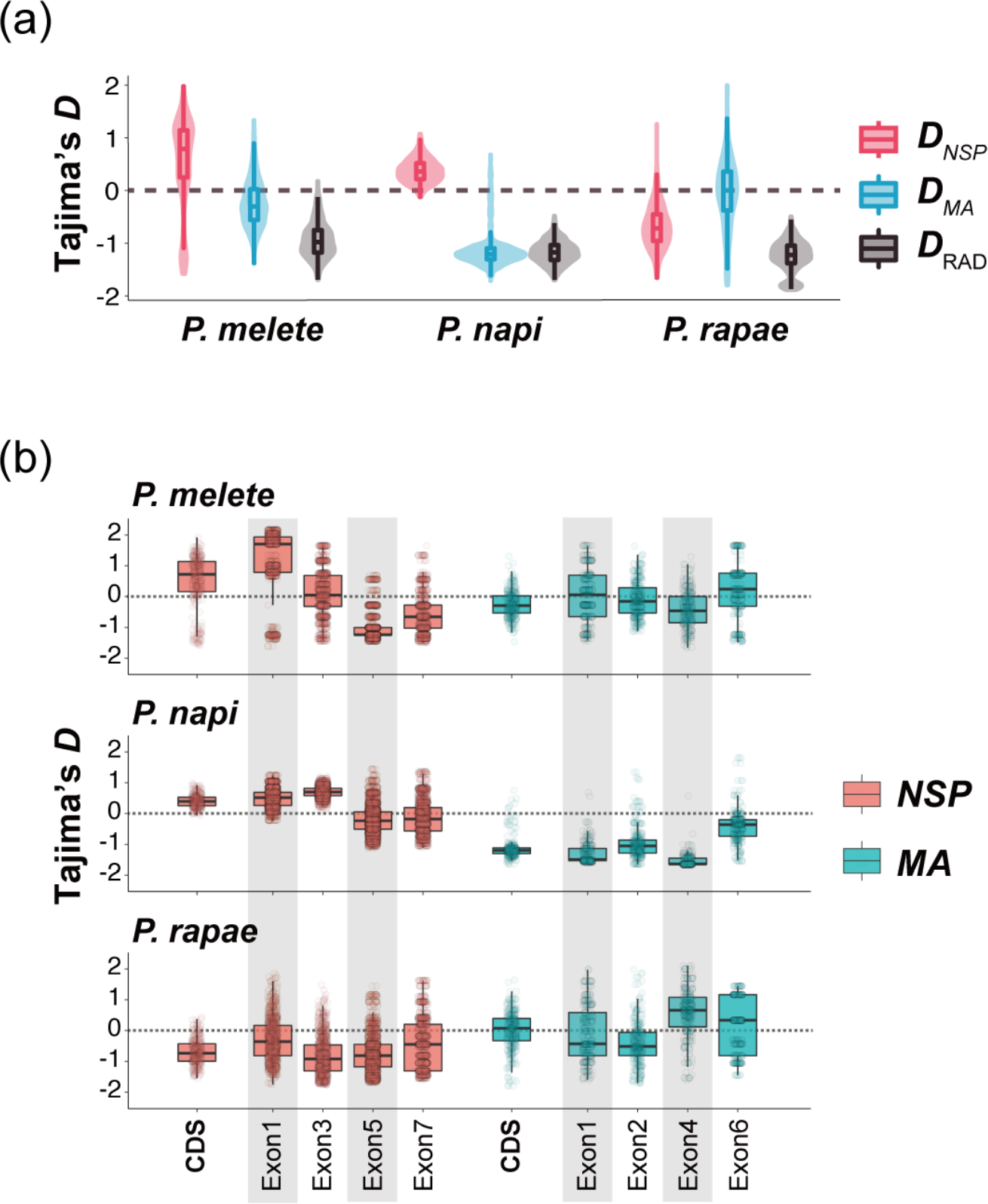
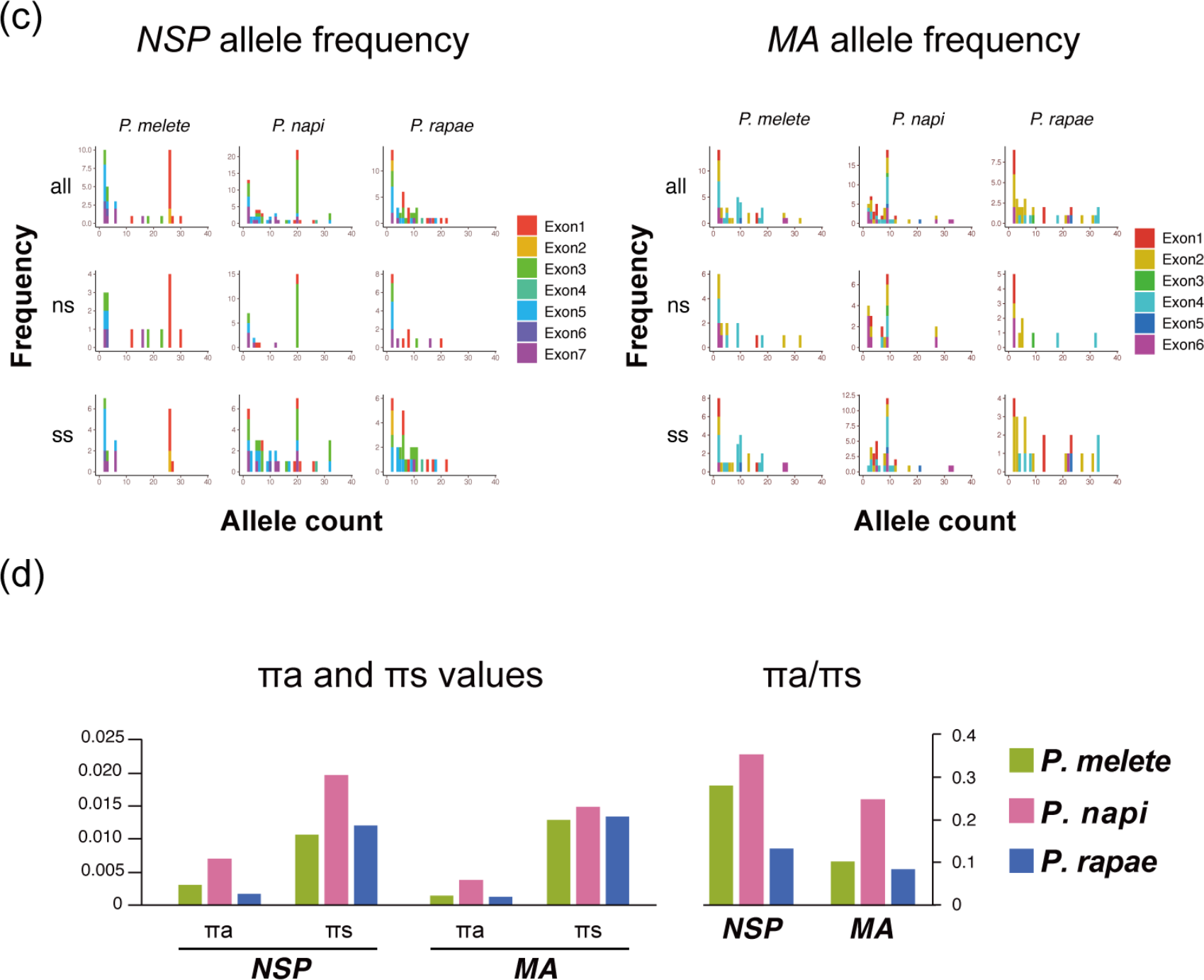
(a) Species-wide distributions of Tajima’s *D* for *NSP*, *MA*, and genome-wide data (RAD). Note high values of *D* for *NSP* in *P. melete* and *P. napi.* (b) Exon-based Tajima’s *D* distribution for *NSP* and *MA*. Only exons > 200 bp are shown. (c) Allele frequency spectra of *NSP* and *MA* based on exon color coding. Exon 1 of *P. melete* and exon 3 of *P. napi NSP*s accumulate positions with intermediate allele frequencies. (d) Non-synonymous (пa) and synonymous nucleotide diversity (пs) at the species level and ratio of non-synonymous to synonymous diversity (пa/пs) of *NSP* and *MA* in three *Pieris* species.

In addition, we compared the ratio of non-synonymous to synonymous nucleotide diversity (пa/пs) of *NSP* and *MA* of the three *Pieris* spp. *MA* had a lower пa/пs ratio compared to *NSP* in all three species (Fig. 5d), suggesting that *MA* was more evolutionary conserved than *NSP* across *Pieris* spp.

#### Divergent selection of *NSP* alleles in *P. napi*

We found that *P. melete* and *P. napi NSP*s, as well as *P. napi MA*s, were all grouped into two clearly distinct clusters (Fig. 3a). Therefore, we tested for potential divergent selection between major alleles, using a population-based diversity test (McDonald & Kreitman, 1991). Results for *P. melete NSP* and *P. napi MA* were non-significant (Table 3), but *P. napi NSP* showed a strong signal of divergent selection (NI = 0.15, *Χ*^2^ = 10.6, *P* = 0.001), caused by an elevated level of non-synonymous *versus* synonymous substitutions between populations from Hokkaido and populations outside of Hokkaido. In order to identify where mutations had occurred, we compared *NSP*s of *P. napi* and *P. melete*, which allowed us to infer the direction of evolutionary change. To our surprise, all non-synonymous substitutions that differentiated the Hokkaido populations of *P. napi* from those of central and southern Japan had occurred outside of Hokkaido, while each two synonymous substitutions had occurred in the lineages leading to the Hokkaido and non-Hokkaido populations. Furthermore, ten non-synonymous and 24 synonymous polymorphisms were specific for populations from central and southern Japan, and only two non-synonymous polymorphisms were specific for populations from Hokkaido (Fig. 6). These patterns of polymorphism indicate that the lineage leading to the extant *P. napi* non-Hokkaido populations experienced strong directional selection on NSP function, followed by stabilizing selection under relaxed constraints, whereas the lineage leading to the extant *P. napi* Hokkaido populations experienced only stabilizing selection.

**Fig. 6.**
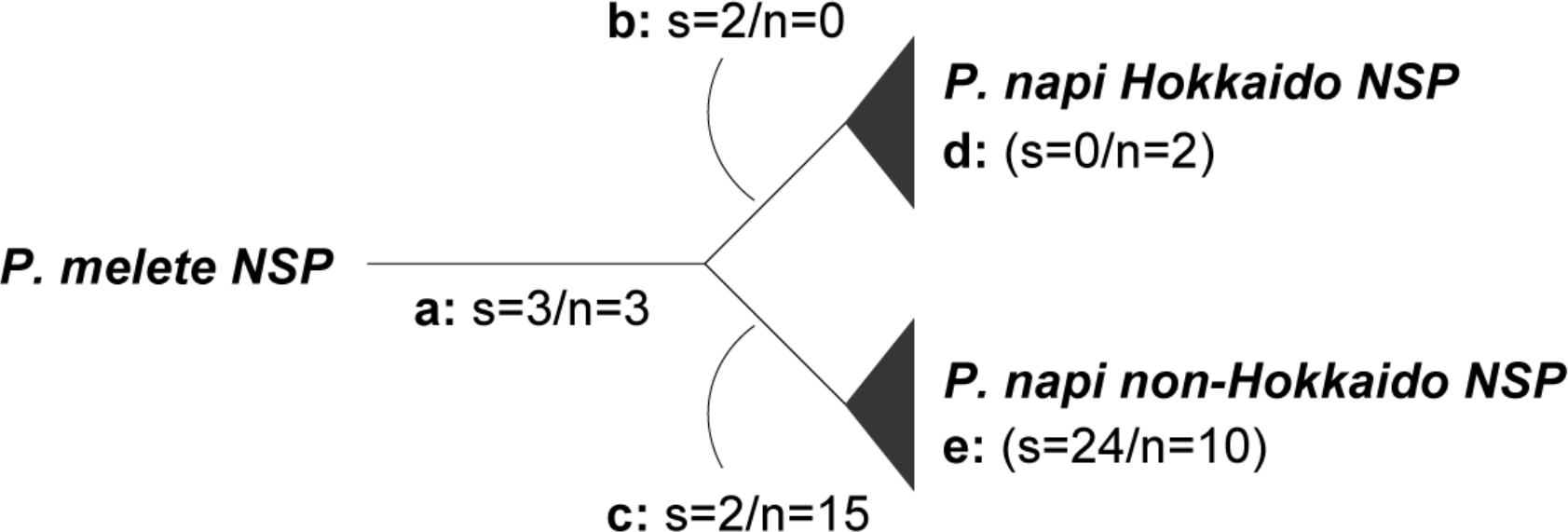
Divergent selection on *NSP* in populations of *Pieris napi*. Synonymous (s) and non-synonymous (n) mutations in *NSP*s on branches of a simplified molecular phylogeny consisting of two *Pieris napi* populations and *Pieris melete*. Polymorphisms segregating in each population are shown in brackets. A very high number of non-synonymous mutations on the non-Hokkaido branch of *Pieris napi* (branch c) indicates positive selection in the lineage leading to the non-Hokkaido populations of *P. napi*. In comparison, *NSP* of the Hokkaido populations appears to be highly conserved.

## Discussion

Several insect counteradaptations against host plant defenses have been characterized at the molecular level (Berenbaum et al., 1996; Gloss et al., 2014; Heidel-Fischer et al., 2019; Li, Schuler, & Berenbaum, 2003; Ratzka et al., 2002; Wittstock et al., 2004) but only few have been investigated for genetic variation in response to host plant usage. Bono et al. (2008) found some evidence for adaptive evolution of two cytochrome P450 genes from the fly *Drosophila mettleri* that were putatively involved in the detoxification of alkaloids from two cactus species. Heidel-Fischer et al. (2010) investigated genetic variation in the *NSP* gene of three European and one US American population of *P. rapae* and found high amounts of amino acid diversity but little evidence for local adaptation. Here, we investigated two genes that underlie the counteradaptation of Pierid butterflies against the GLS-myrosinase defense system of their host plants for potential microevolutionary dynamics in response to host plant usage along a latitudinal cline in Japan. In addition to the crop-dependent *P. rapae*, we included two species that feed on wild Brassicaceae, *P. melete* and *P. napi*. As in the previous study of Heidel-Fischer et al. (2010), we found no evidence for adaptation of *P. rapae NSP*-like genes to local communities of wild Brassicaceae. In contrast, both *P. melete* and *P. napi* showed a significant and positive correlation between the diversity of the *NSP* (but not the *MA*) gene product and host plant diversity (Fig. 4). Both species harbored two distinct *NSP* alleles, which shaped the observed correlations in a species-specific manner.

Hokkaido populations of *P. napi* had a major *NSP* allele that was distinct from alleles found outside of Hokkaido (Fig. 3a). This difference mirrors genome-wide differentiation between both populations (Fig. 2), which suggests two independent colonization events by *P. napi*, one in the north and one in the south of Japan. Differences in the number of acquired polymorphisms in both groups of populations suggest that colonization in the south happened earlier than colonization in the north. Based on the analysis of the mitochondrial *ND5* gene, a recent study concluded that *P. napi* populations on the island of Hokkaido fell into a (south-) western and a (north-) eastern group of populations (Shinkawa et al. 2003), with a hybridization zone in between. Indeed, both Hokkaido populations had elevated genome-wide diversity (Table 1a), and both populations (and in particular Rusutsu) showed signs of gene flow from the south (Fig. 2). Furthermore, *MA* alleles prevalent in the south had apparently replaced the original *MA* alleles in Rusutsu and partially also those in Yubari (Fig. 3d). However, despite gene flow and in contrast to the replacement of *MA* alleles, both Hokkaido populations retained their original *NSP* allele (Fig. 3d). Furthermore, *NSPs* from Hokkaido and from central and southern Japan differed by more non-synonymous substitutions than expected (Table 3), indicative of diversifying selection. However, all amino acid changes that distinguished Hokkaido and non-Hokkaido populations occurred in non-Hokkaido lineage (Fig. 6), indicating that strong directional selection happened in this group, potentially as an adaptive response to novel sets of glucosinolate profiles after colonization. In contrast, both Hokkaido populations displayed very low *NSP* nucleotide diversity, suggesting strong conservation of NSP function. This finding is somewhat surprising because *Rorippa sylvestris*, the preferred host plant species of *P. napi* on the island of Hokkaido (Table S4), was introduced only about 50 years ago (Ohsaki, Ohata, Sato, & Rausher, 2020). It would therefore be helpful to characterize the functional properties of *NSP* gene products from both metapopulations to better understand the contrasting patterns of selection on *NSP* of Japanese *P. napi*.

In contrast to *P. napi*, *P. melete* did not show any indication of population substructure (Fig. 2). In this species, NSP diversity correlated with host plant diversity, which varied along a latitudinal gradient (Fig. 4). Two distinct NSP alleles were present in areas of high host plant diversity, *i.e.*, in central Japan, whereas areas of low host plant diversity were biased towards one of the two alleles, up to fixation of one allele in the Hokkaido populations (Fig. 3d). Although we did not find evidence for diversifying selection, Tajima’s *D* was substantially higher for *NSP* than for *MA* or genome-wide sequence, in particular in the first exon (Fig. 5b). This region indeed harbored an elevated number of linked synonymous and non-synonymous SNPs at intermediate frequency (Fig. 5c), suggesting balancing selection acting on this exon across Japan. Balancing selection is an important mechanism to maintain polymorphisms in natural populations, helping organisms to adapt to environments varying in space and time (Hedrick, 2006). Considering the heterogeneous chemical interactions between herbivores and their host plants in the field, it is not surprising to see such polymorphisms maintained in genes involved in these interactions within natural populations. Indeed, there is evidence on the plant side for balancing selection on genes causing variation in chemical defense (Carley et al., 2021; Kroymann, Donnerhacke, Schnabelrauch, & Mitchell-Olds, 2003; Mitchell- Olds & Schmitt, 2006). By contrast, such patterns have previously not been described for counteradaptive traits of herbivorous insects, mainly due to a lack of information about the genetic bases of such traits. Therefore, *NSP* in Japanese *P. melete* may represent the first example for balancing selection acting on an insect counteradaptation against host plant defenses. Of course, functional data would be helpful to substantiate this idea.

For MA we did not find a clear evolutionary trend. *P. napi* had two distinct MA alleles but these did not show any indication for divergent or balancing selection (Tables 2, 3, Figs. 4, 5). Instead, the comparison of non-synonymous *versus* synonymous diversity suggested that *MA*s were more strongly conserved than *NSP*s in all three *Pieris* species (Fig. 5d). This is consistent with a previous study (Okamura, Sato, Tsuzuki, Murakami, et al., 2019), which found evidence for positive selection only for *NSP* but not for *MA* among different *Pieris* species. Although the functional differences between *NSP* and *MA* are still unknown, both genes displayed distinct expression patterns in response to different GLSs (Okamura, Sato, Tsuzuki, Sawada, et al., 2019). This could indicate that these ancient homologs play different roles in GLS detoxification, with the more evolutionary conserved MAs detoxifying more common GLSs, and NSP acting on rarer GLSs.

Remarkably, the same genes show substantial differences in polymorphism patterns and evolutionary trajectories in three different *Pieris* species that populate Japan. This finding can be partially (but not entirely) explained by different host plant usage. The observed different evolutionary dynamics between *NSP* and its paralog *MA* shows that gene duplication and potential functional differentiation appears to be important for adaptation to chemically heterogeneous host plants in the field. Demographic history, and in particular timing and number of colonization events may also play a major role. Therefore, it would be interesting to investigate the same genes, *NSP* and *MA*, in Pierids collected along a latitudinal cline from the eastern coasts of the Asian continent for comparison.

## Acknowledgements

We thank Domenica Schnabelrauch for her help in sequencing. We are grateful to the members of the community ecology laboratory of Chiba University and Ryo Maeda for their help with field work. We also thank Hanna Heidel-Fischer for comments and suggestions. We thank Emily Wheeler, Boston, for editorial assistance. This work was supported by JSPS KAKENHI Grant-in-Aid for JSPS Fellowships 15J00320 and JSPS Overseas Research Fellowships 202060676 for YO, and Max-Planck-Gesellschaft.

## Data Availability Statement

The RAD-seq short read data have been deposited in the EBI short read archive (SRA) with the following project URL: http://www.ebi.ac.uk/ena/data/view/PRJEB47778.

## Author Contributions

Y.O., A.S., L.K., and A.J.N. carried out the laboratory work. Y.O., M.M., H.V., and J.K. conceived, designed and coordinated the study. Y.O., H.V. and J.K. wrote the manuscript. All authors, drafted parts of the manuscript, gave approval for publication and agree to be accountable for the content.

## Supplementary Tables and Figures

**Table S1.** 19 climate variables used for Maxent

| Variables | Description |
| --- | --- |
| BIO1 | Annual Mean Temperature |
| BIO2 | Mean Diurnal Range (Mean of monthly (max temp - min temp)) |
| BIO3 | Isothermality (BIO2/BIO7) ($\times 100$) |
| BIO4 | Temperature Seasonality (standard deviation $\times 100$) |
| BIO5 | Max Temperature of Warmest Month |
| BIO6 | Min Temperature of Coldest Month |
| BIO7 | Temperature Annual Range (BIO5-BIO6) |
| BIO8 | Mean Temperature of Wettest Quarter |
| BIO9 | Mean Temperature of Driest Quarter |
| BIO10 | Mean Temperature of Warmest Quarter |
| BIO11 | Mean Temperature of Coldest Quarter |
| BIO12 | Annual Precipitation |
| BIO13 | Precipitation of Wettest Month |
| BIO14 | Precipitation of Driest Month |
| BIO15 | Precipitation Seasonality (Coefficient of Variation) |
| BIO16 | Precipitation of Wettest Quarter |
| BIO17 | Precipitation of Driest Quarter |
| BIO18 | Precipitation of Warmest Quarter |
| BIO19 | Precipitation of Coldest Quarter |

**Table S2.** Sampling sites and observed number of Brassicaceae

| Sampling Sites | Yubari | Rusutsu | Oshamanbe | Fukushima | Nagano | Chiba | Kanagawa | Nara | Tokushima | Miyazaki | Yonaguni |
| --- | --- | --- | --- | --- | --- | --- | --- | --- | --- | --- | --- |
| latitude | 43.0 | 42.7 | 42.6 | 37.8 | 36.3 | 35.5 | 35.5 | 34.3 | 33.9 | 32.4 | 24.5 |
| longitude | 142.1 | 140.8 | 140.3 | 140.6 | 137.8 | 140.2 | 139.2 | 136.0 | 134.1 | 131.2 | 123.0 |
| Plant diversity | 1.0224 | 0.7115 | 0.6374 | 1.6662 | 2.1451 | 1.7968 | 1.3765 | 1.8850 | 1.5404 | 0.8650 | 0.3125 |
| **Plant Species** |  |  |  |  |  |  |  |  |  |  |  |
| *Arabidopsis thaliana* | 0 | 0 | 0 | 0 | 60 | 20 | 0 | 0 | 0 | 0 | 0 |
| *Arabidopsis halleri* | 0 | 0 | 0 | 0 | 0 | 0 | 0 | 83 | 0 | 0 | 0 |
| *Arabidopsis kamchatica* | 0 | 0 | 0 | 0 | 46 | 0 | 0 | 0 | 0 | 0 | 0 |
| *Arabis flagellosa* | 0 | 0 | 0 | 0 | 0 | 0 | 0 | 150 | 0 | 245 | 0 |
| *Arabis hirsuta* | 0 | 0 | 0 | 78 | 0 | 0 | 72 | 12 | 12 | 0 | 0 |
| *Barbarea orthoceras* | 6 | 0 | 15 | 7 | 72 | 0 | 0 | 0 | 4 | 0 | 0 |
| *Brassica napus* | 0 | 0 | 0 | 0 | 51 | 45 | 0 | 10 | 3 | 0 | 0 |
| *Cardamine impatiens* | 0 | 0 | 0 | 0 | 12 | 4 | 7 | 0 | 5 | 0 | 0 |
| *Cardamine kiusiana* | 0 | 0 | 0 | 0 | 0 | 0 | 0 | 0 | 14 | 0 | 0 |
| *Cardamine leucantha* | 438 | 20 | 130 | 42 | 140 | 0 | 0 | 0 | 0 | 0 | 0 |
| *Cardamine regeliana* | 0 | 0 | 0 | 0 | 81 | 125 | 0 | 0 | 0 | 0 | 0 |
| *Cardamine occulta* | 30 | 0 | 12 | 45 | 210 | 60 | 18 | 31 | 0 | 13 | 25 |
| *Draba nemorosa* | 0 | 0 | 0 | 0 | 140 | 0 | 0 | 0 | 0 | 0 | 0 |
| *Eutrema japonicum* | 0 | 0 | 0 | 130 | 15 | 0 | 30 | 0 | 0 | 0 | 0 |
| *Eutrema tenue* | 47 | 0 | 0 | 0 | 0 | 0 | 0 | 0 | 0 | 0 | 0 |
| *Lepidium virginicum* | 0 | 0 | 0 | 0 | 0 | 0 | 0 | 10 | 0 | 0 | 240 |
| *Nasturtium officinale* | 0 | 0 | 0 | 23 | 0 | 200 | 0 | 174 | 60 | 0 | 0 |
| *Orychophragmus violaceus* | 0 | 0 | 0 | 0 | 0 | 140 | 0 | 0 | 0 | 0 | 0 |
| *Raphanus sativus* | 0 | 0 | 0 | 0 | 8 | 0 | 0 | 0 | 0 | 0 | 0 |
| *Rorippa dubia* | 0 | 0 | 0 | 0 | 0 | 0 | 0 | 62 | 0 | 58 | 0 |
| *Rorippa indica* | 0 | 0 | 2 | 25 | 0 | 40 | 58 | 75 | 70 | 27 | 0 |
| *Rorippa palustris* | 0 | 101 | 0 | 0 | 21 | 10 | 0 | 10 | 0 | 0 | 0 |
| *Rorippa sylvestris* | 262 | 317 | 0 | 0 | 0 | 0 | 0 | 0 | 0 | 0 | 0 |
| *Turritis glabra* | 0 | 0 | 0 | 0 | 0 | 0 | 0 | 0 | 30 | 0 | 0 |

**Table S3.**
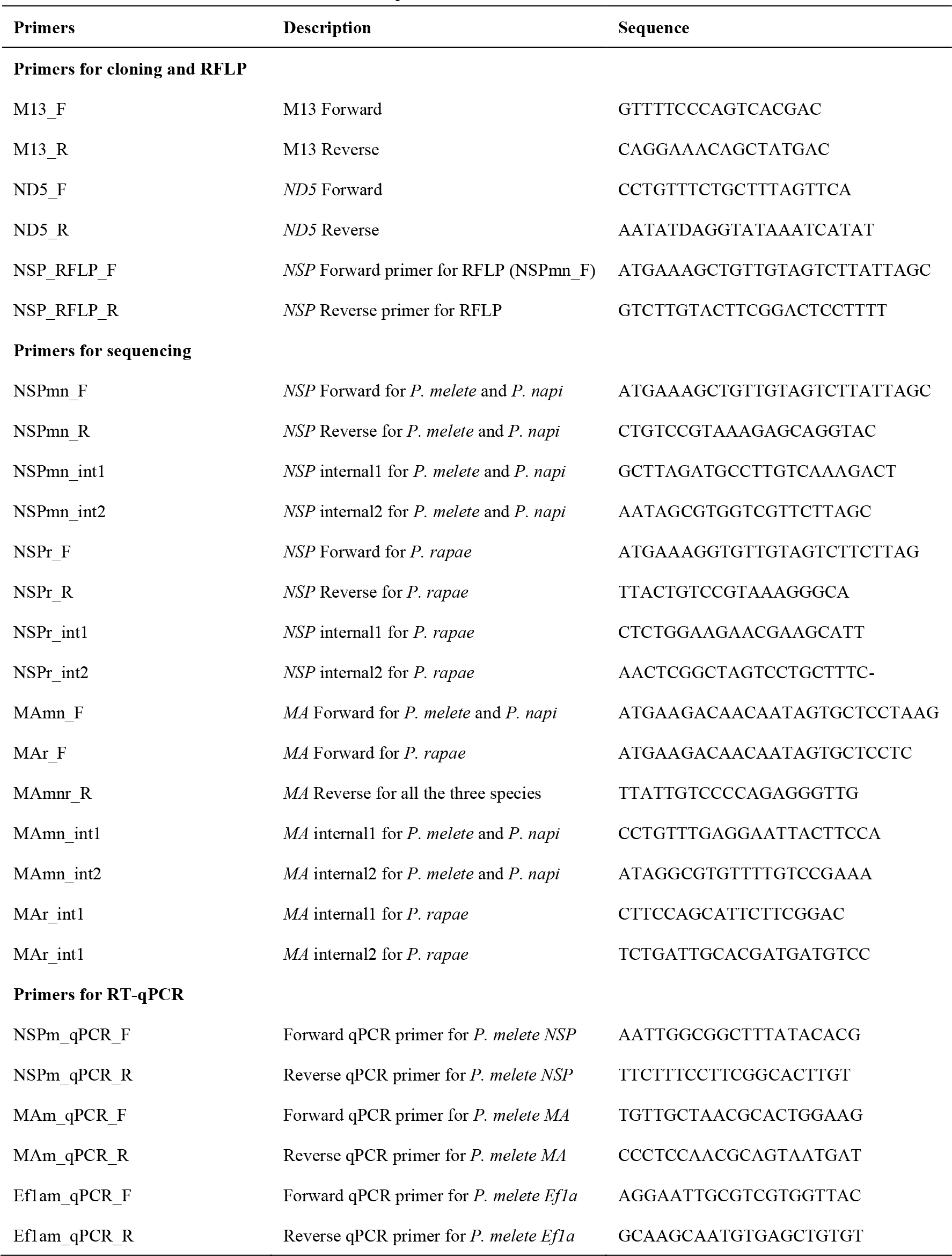
Primers used in this study

**Table S4.** Host plant usage patterns of three *Pieris* species at different sampling sites Columns show ‘observed number of larvae’ and ‘checked number of plants’. Hyphens (-) indicate absence of a plant species at the sampling site.

| *P. melete* |  |  |  |  |  |  |  |  |  |  |  |  |
| --- | --- | --- | --- | --- | --- | --- | --- | --- | --- | --- | --- | --- |
| Plant sp./Sampling Sites | Yubari | Rusutsu | Oshamanbe | Fukushima | Nagano | Chiba | Kanagawa | Nara | Tokushima | Miyazaki | Yonaguni | SUM |
| *Arabidopsis halleri* | - | - | - | - | - | - | - | 0/83 | - | - | - | 0/83 |
| *Arabidopsis kamchatica* | - | - | - | - | 0/46 | - | - | - | - | - | - | 0/46 |
| *Arabidopsis thaliana* | - | - | - | - | 0/60 | 0/20 | - | - | - | - | - | 0/80 |
| *Arabis flagellosa* | - | - | - | - | - | - | - | 0/150 | - | 0/245 | - | 0/395 |
| *Arabis hirsuta* | - | - | - | 27/78 | - | - | 10/72 | 0/12 | 28/12 | - | - | 65/174 |
| *Barbarea orthoceras* | 0/6 | - | 0/15 | 0/7 | 0/72 | - | - | - | 0/4 | - | - | 0/104 |
| *Brassica napus* | - | - | - | - | 0/51 | 0/45 | - | 0/10 | 0/3 | - | - | 0/109 |
| *Cardamine impatiens* | - | - | - | - | 0/12 | 0/4 | 9/7 | - | 6/5 | - | - | 15/28 |
| *Cardamine kiusiana* | - | - | - | - | - | - | - | - | 10/14 | - | - | 10/14 |
| *Cardamine leucantha* | 19/438 | 0/20 | 24/130 | 16/42 | 35/140 | - | - | - | - | - | - | 94/770 |
| *Cardamine occulta* | 0/30 | - | 0/12 | 4/45 | 0/210 | 0/60 | 0/18 | 2/31 | - | 0/13 | 0/25 | 6/444 |
| *Cardamine regeliana* | - | - | - | - | 0/81 | 0/125 | - | - | - | - | - | 0/206 |
| *Draba nemorosa* | - | - | - | - | 0/140 | - | - | - | - | - | - | 0/140 |
| *Eutrema japonicum* | - | - | - | 0/130 | 0/15 | - | 0/30 | - | - | - | - | 0/175 |
| *Eutrema tenue* | 0/47 | - | - | - | - | - | - | - | - | - | - | 0/47 |
| *Lepidium virginicum* | - | - | - | - | - | - | - | 0/10 | - | - | 0/240 | 0/250 |
| *Nasturtium officinale* | - | - | - | 0/23 | - | 6/200 | - | 0/174 | 0/6 | - | - | 6/403 |
| *Orychophragmus violaceus* | - | - | - | - | - | 88/140 | - | - | - | - | - | 88/140 |
| *Raphanus sativus* | - | - | - | - | 0/8 | - | - | - | - | - | - | 0/8 |
| *Rorippa dubia* | - | - | - | - | - | - | - | 1/62 | - | 0/58 | - | 1/120 |
| *Rorippa indica* | - | - | 0/2 | 0/25 | - | 18/40 | 25/58 | 48/75 | 85/70 | 0/27 | - | 176/297 |
| *Rorippa palustris* | - | 0/101 | - | - | 0/21 | 0/10 | - | 0/10 | - | - | - | 0/142 |
| *Rorippa sylvestris* | 1/262 | 0/317 | - | - | - | - | - | - | - | - | - | 1/579 |
| *Turritis glabra* | - | - | - | - | - | - | - | - | 21/30 | - | - | 21/30 |
| **SUM** | 20/783 | 0/438 | 24/159 | 47/350 | 35/856 | 112/644 | 44/185 | 51/617 | 150/144 | 0/343 | 0/265 | 483/4784 |

***P. napi***
| Plant sp./Sampling Sites | Yubari | Rusutsu | Oshamanbe | Fukushima | Nagano | Chiba | Kanagawa | Nara | Tokushima | Miyazaki | Yonaguni | **SUM** |
| --- | --- | --- | --- | --- | --- | --- | --- | --- | --- | --- | --- | --- |
| *Arabidopsis halleri* | - | - | - | - | - | - | - | 11/83 | - | - | - | 11/83 |
| *Arabidopsis kamchatica* | - | - | - | - | 7/46 | - | - | - | - | - | - | 7/46 |
| *Arabidopsis thaliana* | - | - | - | - | 0/60 | 0/20 | - | - | - | - | - | 0/80 |
| *Arabis flagellosa* | - | - | - | - | - | - | - | 10/150 | - | 34/245 | - | 44/395 |
| *Arabis hirsuta* | - | - | - | 19/78 | - | - | 7/72 | 5/12 | 0/12 | - | - | 31/174 |
| *Barbarea orthoceras* | 0/6 | - | 0/15 | 0/7 | 29/72 | - | - | - | 0/4 | - | - | 29/104 |
| *Brassica napus* | - | - | - | - | 0/51 | 0/45 | - | 0/10 | 0/3 | - | - | 0/109 |
| *Cardamine impatiens* | - | - | - | - | 0/12 | 0/4 | 2/7 | - | 0/5 | - | - | 2/28 |
| *Cardamine kiusiana* | - | - | - | - | - | - | - | - | 0/14 | - | - | 0/14 |
| *Cardamine leucantha* | 5/438 | 0/20 | 0/130 | 4/42 | 0/140 | - | - | - | - | - | - | 9/770 |
| *Cardamine occulta* | 1/30 | - | 0/12 | 0/45 | 39/210 | 0/60 | 0/18 | 4/31 | - | 0/13 | 0/25 | 44/444 |
| *Cardamine regeliana* | - | - | - | - | 2/81 | 0/125 | - | - | - | - | - | 2/206 |
| *Draba nemorosa* | - | - | - | - | 0/140 | - | - | - | - | - | - | 0/140 |
| *Eutrema japonicum* | - | - | - | 0/130 | 0/15 | - | 0/30 | - | - | - | - | 0/175 |
| *Eutrema tenue* | 1/47 | - | - | - | - | - | - | - | - | - | - | 1/47 |
| *Lepidium virginicum* | - | - | - | - | - | - | - | 0/10 | - | - | 0/240 | 0/250 |
| *Nasturtium officinale* | - | - | - | 0/23 | - | 0/200 | - | 0/174 | 0/6 | - | - | 0/403 |
| *Orychophragmus violaceus* | - | - | - | - | - | 0/140 | - | - | - | - | - | 0/140 |
| *Raphanus sativus* | - | - | - | - | 0/8 | - | - | - | - | - | - | 0/8 |
| *Rorippa dubia* | - | - | - | - | - | - | - | 3/62 | - | 0/58 | - | 3/120 |
| *Rorippa indica* | - | - | 0/2 | 1/25 | - | 0/40 | 5/58 | 0/75 | 0/70 | 0/27 | - | 6/297 |
| *Rorippa palustris* | - | 6/101 | - | - | 0/21 | 0/10 | - | 0/10 | - | - | - | 6/142 |
| *Rorippa sylvestris* | 21/262 | 37/317 | - | - | - | - | - | - | - | - | - | 58/579 |
| *Turritis glabra* | - | - | - | - | - | - | - | - | 0/30 | - | - | 0/30 |
| **SUM** | 28/783 | 43/438 | 0/159 | 24/350 | 77/856 | 0/644 | 14/185 | 33/617 | 0/144 | 34/343 | 0/265 | 253/4784 |

| ***P. rapae*** |  |  |  |  |  |  |  |  |  |  |  |  |
| --- | --- | --- | --- | --- | --- | --- | --- | --- | --- | --- | --- | --- |
| Plant sp./Sampling Sites | Yubari | Rusutsu | Oshamanbe | Fukushima | Nagano | Chiba | Kanagawa | Nara | Tokushima | Miyazaki | Yonaguni | SUM |
| *Arabidopsis halleri* | - | - | - | - | - | - | - | 0/83 | - | - | - | 0 /83 |
| *Arabidopsis kamchatica* | - | - | - | - | 0/46 | - | - | - | - | - | - | 0 /46 |
| *Arabidopsis thaliana* | - | - | - | - | 0/60 | 0/20 | - | - | - | - | - | 0 /80 |
| *Arabis flagellosa* | - | - | - | - | - | - | - | 0/150 | - | 0/245 | - | 0 /395 |
| *Arabis hirsuta* | - | - | - | 0/78 | - | - | 0/72 | 0/12 | 0/12 | - | - | 0 /174 |
| *Barbarea orthoceras* | 2/6 | - | 0/15 | 0/7 | 0/72 | - | - | - | 0/4 | - | - | 2 /104 |
| *Brassica napus* | - | - | - | - | 35/51 | 34/45 | - | 0/10 | 0/3 | - | - | 69 /109 |
| *Cardamine impatiens* | - | - | - | - | 0/12 | 0/4 | 0/7 | - | 0/5 | - | - | 0 /28 |
| *Cardamine kiusiana* | - | - | - | - | - | - | - | - | 25/14 | - | - | 25 /14 |
| *Cardamine leucantha* | 0/438 | 0/20 | 0/130 | 0/42 | 0/140 | - | - | - | - | - | - | 0 /770 |
| *Cardamine occulta* | 0/30 | - | 0/12 | 0/45 | 0/210 | 0/60 | 0/18 | 0/31 | - | 0/13 | 0/25 | 0 /444 |
| *Cardamine regeliana* | - | - | - | - | 0/81 | 0/125 | - | - | - | - | - | 0 /206 |
| *Draba nemorosa* | - | - | - | - | 0/140 | - | - | - | - | - | - | 0 /140 |
| *Eutrema japonicum* | - | - | - | 0/130 | 0/15 | - | 0/30 | - | - | - | - | 0 /175 |
| *Eutrema tenue* | 0/47 | - | - | - | - | - | - | - | - | - | - | 0 /47 |
| *Lepidium virginicum* | - | - | - | - | - | - | - | 0/10 | - | - | 8/240 | 8 /250 |
| *Nasturtium officinale* | - | - | - | 1/23 | - | 0/200 | - | 13/174 | 6/6 | - | - | 20 /403 |
| *Orychophragmus violaceus* | - | - | - | - | - | 0/140 | - | - | - | - | - | 0 /140 |
| *Raphanus sativus* | - | - | - | - | 4/8 | - | - | - | - | - | - | 4 /8 |
| *Rorippa dubia* | - | - | - | - | - | - | - | 0/62 | - | 0/58 | - | 0 /120 |
| *Rorippa indica* | - | - | 0/2 | 11/25 | - | 0/40 | 0/58 | 7/75 | 13/70 | 0/27 | - | 31 /297 |
| *Rorippa palustris* | - | 20/101 | - | - | 0/21 | 0/10 | - | 0/10 | - | - | - | 20 /142 |
| *Rorippa sylvestris* | 29/262 | 1/317 | - | - | - | - | - | - | - | - | - | 30 /579 |
| *Turritis glabra* | - | - | - | - | - | - | - | - | 0/30 | - | - | 0 /30 |
| **SUM** | 31 /783 | 21 /438 | 0 /159 | 12 /350 | 39 /856 | 34 /644 | 0 /185 | 20 /617 | 44 /144 | 0 /343 | 8 /265 | 209 /4784 |

**Table S5.** Statistics for *NSP* and *MA*

| Species | Sampling sites | Sequences | Number of sites | S sites | *NSP* $\pi$ | AA diversity | Tajima's | *P* value |
| --- | --- | --- | --- | --- | --- | --- | --- | --- |
| *P. melete* | ALL | 69 | 1860 | 34 | 0.0047 | 0.0070 | 0.561 | - |
|  | Yubari (Hokkaido) | 10 | 1860 | 5 | 0.0007 | 0.0009 | -1.035 | 0.188 |
|  | Oshamanbe | 10 | 1860 | 4 | 0.0005 | 0.0003 | -1.245 | 0.116 |
|  | Fukushima | 9 | 1860 | 23 | 0.0052 | 0.0080 | 0.738 | 0.803 |
|  | Nagano | 10 | 1860 | 18 | 0.0047 | 0.0070 | 1.772 | 0.980 |
|  | Chiba | 10 | 1860 | 22 | 0.0045 | 0.0065 | 0.406 | 0.685 |
|  | Nara | 10 | 1857-1860 | 24 | 0.0049 | 0.0069 | 0.314 | 0.662 |
|  | Tokushima | 10 | 1857-1860 | 27 | 0.0059 | 0.0091 | 0.773 | 0.815 |
| *P. napi* | ALL | 69 | 1860 | 66 | 0.0098 | 0.0156 | 0.370 | - |
|  | Yubari (Hokkaido) | 10 | 1860 | 5 | 0.0006 | 0.0015 | -1.388 | 0.094 |
|  | Rusutsu (Hokkaido) | 10 | 1860 | 2 | 0.0004 | 0.0012 | 0.120 | 0.686 |
|  | Fukushima | 9 | 1860 | 27 | 0.0052 | 0.0043 | -0.107 | 0.483 |
|  | Nagano | 10 | 1860 | 29 | 0.0056 | 0.0039 | 0.132 | 0.584 |
|  | Kanagawa | 10 | 1860 | 27 | 0.0053 | 0.0041 | 0.162 | 0.610 |
|  | Nara | 10 | 1860 | 26 | 0.0051 | 0.0043 | 0.178 | 0.614 |
|  | Miyazaki | 10 | 1860 | 19 | 0.0037 | 0.0012 | 0.182 | 0.615 |
| *P. rapae* | ALL | 74 | 1869 | 43 | 0.0039 | 0.0037 | -0.704 | - |
|  | Yubari (Hokkaido) | 10 | 1869 | 22 | 0.0029 | 0.0032 | -1.485 | 0.062 |
|  | Rusutsu (Hokkaido) | 9 | 1869 | 21 | 0.0033 | 0.0030 | -1.031 | 0.164 |
|  | Fukushima | 9 | 1869 | 23 | 0.0047 | 0.0038 | 0.152 | 0.587 |
|  | Nagano | 10 | 1869 | 21 | 0.0029 | 0.0028 | -1.257 | 0.105 |
|  | Chiba | 8 | 1869 | 18 | 0.0033 | 0.0030 | -0.676 | 0.281 |
|  | Nara | 10 | 1869 | 26 | 0.0043 | 0.0038 | -0.664 | 0.265 |
|  | Tokushima | 10 | 1869 | 28 | 0.0053 | 0.0052 | -0.221 | 0.518 |
|  | Yonaguni (Okinawa) | 8 | 1869 | 18 | 0.0036 | 0.0037 | -0.485 | 0.427 |

| Species | Sampling sites | Sequences | Number of sites | S sites | *MA* $\pi$ | AA diversity | Tajima's *D* | *P* value |
| --- | --- | --- | --- | --- | --- | --- | --- | --- |
| *P. melete* | ALL | 68 | 1896 | 44 | 0.0048 | 0.0056 | -0.2495 | - |
|  | Yubari (Hokkaido) | 10 | 1896 | 21 | 0.0035 | 0.0032 | -0.5370 | 0.314 |
|  | Oshamanbe | 10 | 1896 | 23 | 0.0045 | 0.0043 | 0.1832 | 0.617 |
|  | Fukushima | 10 | 1896 | 20 | 0.0044 | 0.0065 | 0.8698 | 0.840 |
|  | Nagano | 10 | 1896 | 24 | 0.0041 | 0.0049 | -0.4326 | 0.348 |
|  | Chiba | 8 | 1896 | 16 | 0.0039 | 0.0031 | 0.9582 | 0.859 |
|  | Nara | 10 | 1896 | 31 | 0.0059 | 0.0076 | 0.0867 | 0.579 |
|  | Tokushima | 10 | 1896 | 29 | 0.0052 | 0.0069 | -0.2318 | 0.437 |
| *P. napi* | ALL | 70 | 1896 | 56 | 0.0061 | 0.0087 | -1.1531 | - |
|  | Yubari (Hokkaido) | 10 | 1896 | 39 | 0.0056 | 0.0088 | -1.2381 | 0.134 |
|  | Rusutsu | 10 | 1896 | 11 | 0.0023 | 0.0022 | 0.4869 | 0.711 |
|  | Fukushima | 10 | 1896 | 17 | 0.0029 | 0.0022 | -0.3511 | 0.389 |
|  | Nagano | 10 | 1896 | 14 | 0.0021 | 0.0021 | -0.9809 | 0.176 |
|  | Kanagawa | 10 | 1896 | 15 | 0.0029 | 0.0029 | 0.2393 | 0.632 |
|  | Nara | 10 | 1896 | 14 | 0.0028 | 0.0031 | 0.3970 | 0.687 |
|  | Miyazaki | 10 | 1896 | 10 | 0.0021 | 0.0020 | 0.5833 | 0.747 |
| *P. rapae* | ALL | 76 | 1896 | 34 | 0.0038 | 0.0025 | -0.0747 | - |
|  | Yubari (Hokkaido) | 10 | 1896 | 18 | 0.0040 | 0.0033 | 0.8915 | 0.844 |
|  | Rusutsu | 9 | 1896 | 20 | 0.0042 | 0.0020 | 0.3914 | 0.687 |
|  | Fukushima | 10 | 1896 | 18 | 0.0033 | 0.0024 | -0.0542 | 0.508 |
|  | Nagano | 10 | 1896 | 15 | 0.0032 | 0.0025 | 0.6447 | 0.772 |
|  | Chiba | 9 | 1896 | 16 | 0.0038 | 0.0025 | 1.0538 | 0.881 |
|  | Nara | 10 | 1896 | 14 | 0.0027 | 0.0016 | 0.2119 | 0.625 |
|  | Tokushima | 10 | 1896 | 22 | 0.0041 | 0.0012 | -0.0399 | 0.513 |
|  | Yonaguni | 8 | 1896 | 22 | 0.0046 | 0.0033 | 0.0754 | 0.548 |

**Fig. S1.**
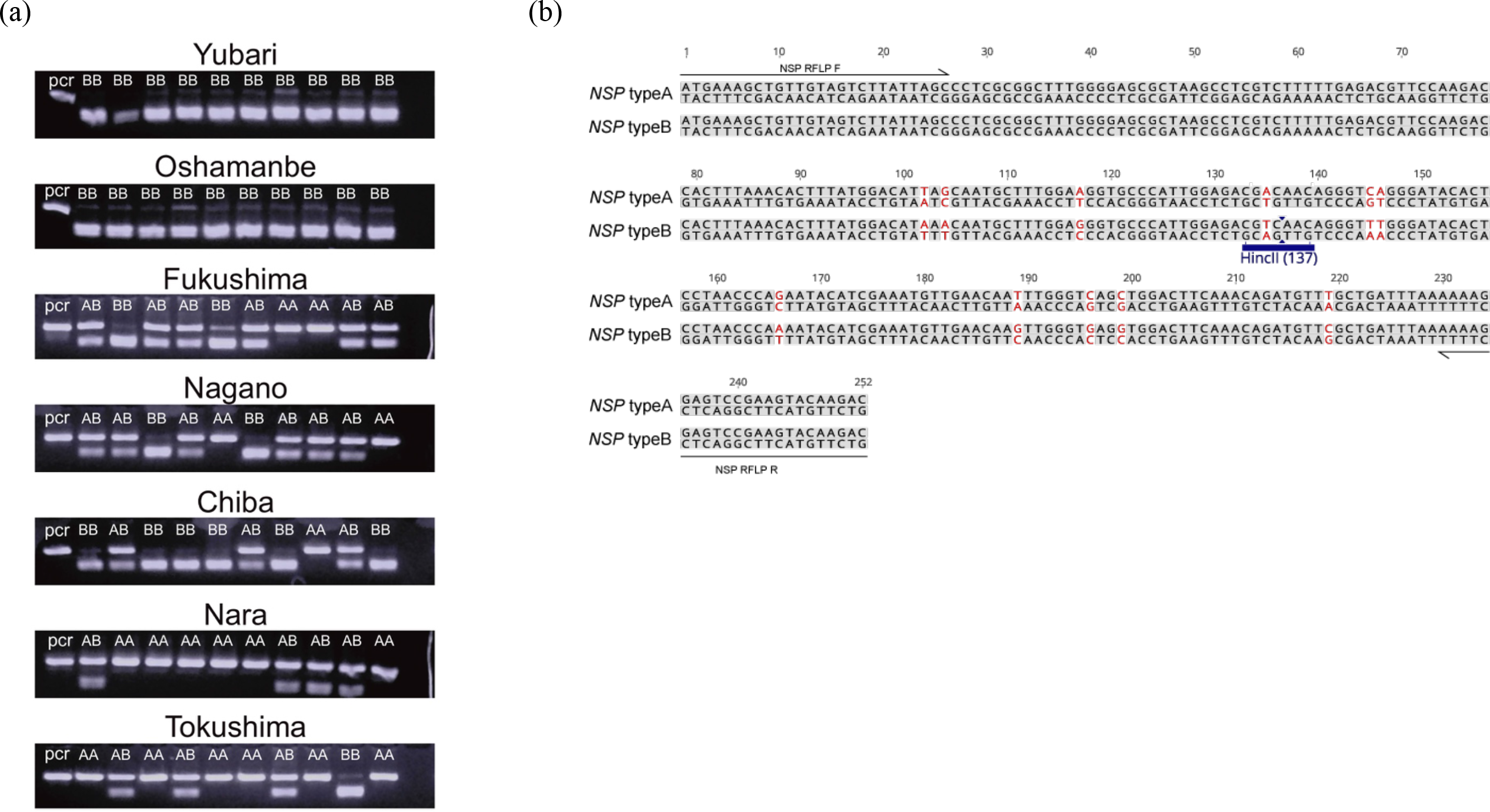
(a) PCR-RFLP of NSP exon 1 from seven populations of P. melete. “pcr” refers to an undigested PCR product, corresponding to ‘type A’. Note that ‘type B’ bands in the agarose gels consist of two restriction fragments of similar size. (b) Exon 1 sequences of type A and type B *NSP* with *Hin*cII restriction site in type B. SNPs distinguishing types A and B of *NSP* are shown in red.

**Fig. S2.**
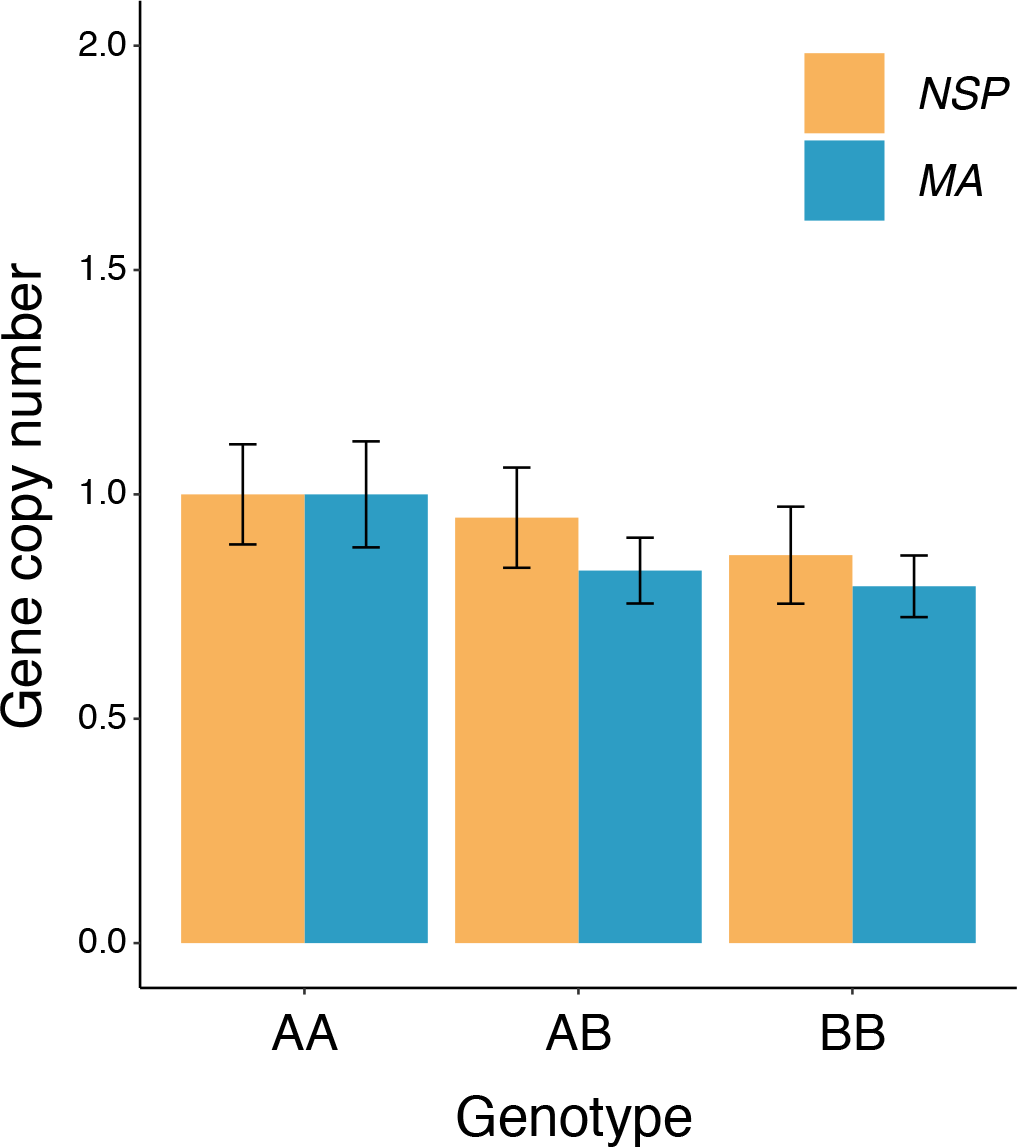
*NSP* and *MA* gene copy numbers in *P. melete*. Genotypes (AA, AB and BB) are the same as in Fig. S1. RT-qPCR was done with N = 8 replicates per genotype and *Ef1α* as a reference gene. Data were standardized with mean of genotype AA set as 1. Bars shows means (± SE). Genotypes do not differ in *NSP* (yellow bars) or *MA* (blue bars) gene copy numbers.

**Fig. S3.**
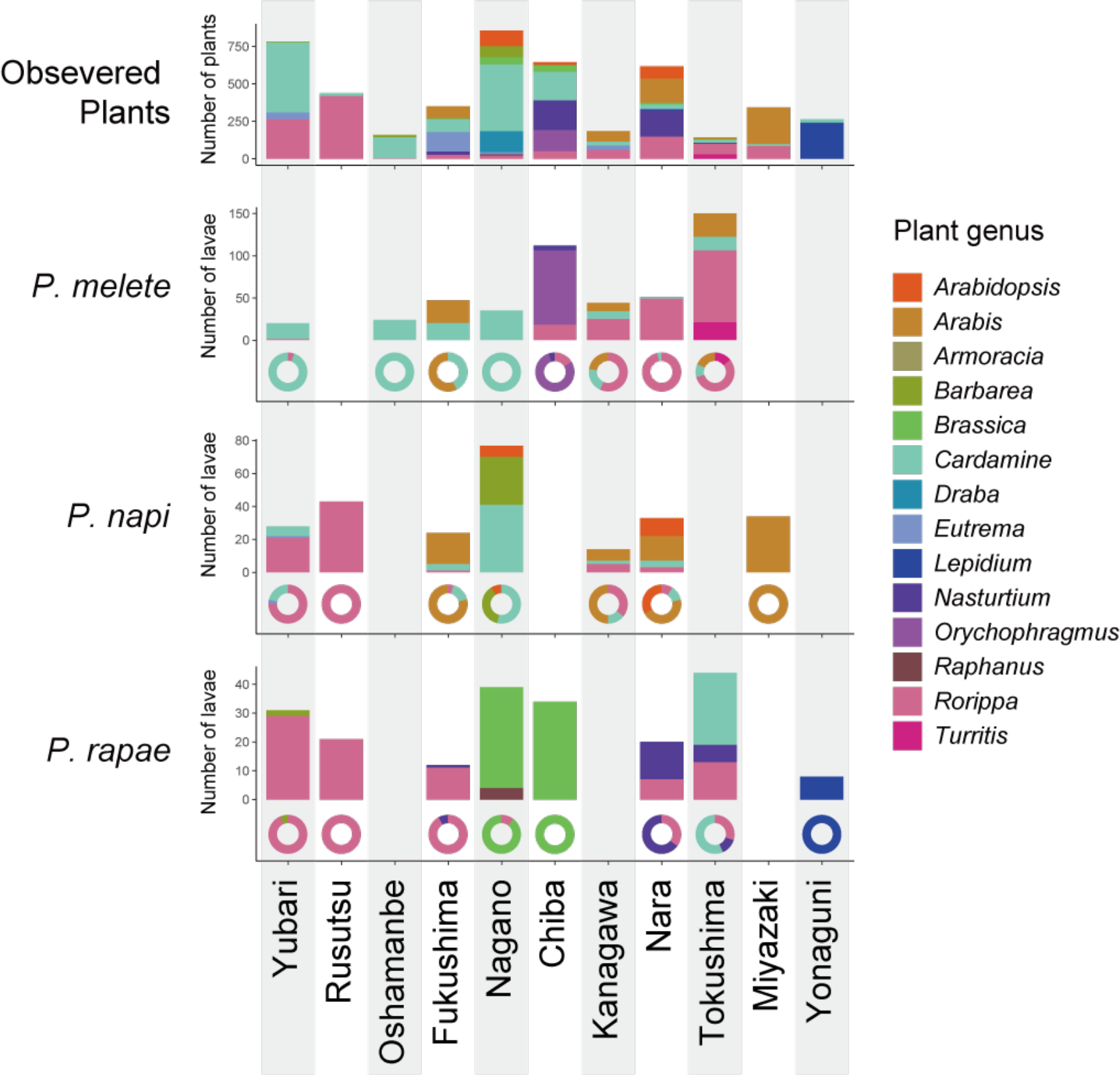
Patterns of host plant use of *Pieris* species in the field. The plant species data were summarized to genus level for visualization. The pie charts show composition of plant genus and each bar show number of individuals (either plants or larvae).

